# DNA damage-induced interaction between a lineage addiction oncogenic transcription factor and the MRN complex shapes a tissue-specific DNA Damage Response and cancer predisposition

**DOI:** 10.1101/2023.04.21.537819

**Authors:** Romuald Binet, Jean-Philippe Lambert, Marketa Tomkova, Samuel Tischfield, Arianna Baggiolini, Sarah Picaud, Sovan Sarkar, Pakavarin Louphrasitthiphol, Diogo Dias, Suzanne Carreira, Timothy Humphrey, Panagis Fillipakopoulos, Richard White, Colin R Goding

## Abstract

Since genome instability can drive cancer initiation and progression, cells have evolved highly effective and ubiquitous DNA Damage Response (DDR) programs. However, some cells, in skin for example, are normally exposed to high levels of DNA damaging agents. Whether such high-risk cells possess lineage-specific mechanisms that tailor DNA repair to the tissue remains largely unknown. Here we show, using melanoma as a model, that the microphthalmia-associated transcription factor MITF, a lineage addition oncogene that coordinates many aspects of melanocyte and melanoma biology, plays a non-transcriptional role in shaping the DDR. On exposure to DNA damaging agents, MITF is phosphorylated by ATM/DNA-PKcs, and unexpectedly its interactome is dramatically remodelled; most transcription (co)factors dissociate, and instead MITF interacts with the MRE11-RAD50-NBS1 (MRN) complex. Consequently, cells with high MITF levels accumulate stalled replication forks, and display defects in homologous recombination-mediated repair associated with impaired MRN recruitment to DNA damage. In agreement, high MITF levels are associated with increased SNV burden in melanoma. Significantly, the SUMOylation-defective MITF-E318K melanoma predisposition mutation recapitulates the effects of ATM/DNA-PKcs-phosphorylated MITF. Our data suggest that a non-transcriptional function of a lineage-restricted transcription factor contributes to a tissue-specialised modulation of the DDR that can impact cancer initiation.

## Introduction

Preserving the integrity of the genome is critical for survival and the faithful inheritance of genetic information following DNA replication. Since the genome is subject to a wide range of insults, including DNA-replication stress, irradiation, and exposure to DNA damaging agents including chemotherapeutic drugs, cells have evolved an arsenal of sophisticated mechanisms to repair DNA damage and maintain genome integrity. Failure of the repair pathways to accurately resolve DNA damage can lead to cell death or disease, including cancer. One of the most detrimental types of DNA damage are double strand breaks (DSBs) that are repaired by two principal mechanisms: non-homologous end joining (NHEJ) and homologous recombination-mediated repair (HRR) (Ceccaldi et al. 2016). In NHEJ, the Ku70/80/DNA-PK complex protects exposed DNA ends before the Ligase IV/XRCC4/XLF complex mediates end re-joining. This process can occur in all phases of the cell cycle but is error-prone and may consequently cause detrimental mutations or deletions. By contrast, HRR relies on an initial resection step controlled by the MRE11/RAD50/NBS1 (MRN) complex and the nuclease CtIP (Buis et al. 2012; Paull 2018). Newly generated 3’ overhangs are then protected by RPA that serves as a platform to recruit the RAD51 recombinase that in turn searches for a template DNA to invade and copy. As a result, HRR is restricted to S and G2 phases of the cell cycle but mediates more faithful repair than NHEJ. Alternative HR-mediated repair mechanisms have also been described, namely single-strand annealing (SSA), alternative end-joining (alt-EJ), break-induced replication (BIR) and RNA-directed DNA repair (RDDR) (Yang and Qi 2015; Bhargava et al. 2016; Ceccaldi et al. 2016; Kramara et al. 2018). Involvement of the HRR and associated pathways depends on the efficiency of end resection, availability of a template DNA, or absence of one end of the DSB and can be independent of RAD51 and involve other recombinases like RAD52 and RAD59 (Rossi et al. 2021). Although DNA damage repair (DDR) mechanisms appear to be largely conserved in different tissues, accumulating evidence suggests that their function might be modulated by tissue-restricted factors that would tailor repair pathways to reflect the demands of specific cell types (Chao and Lipkin 2006; D’Errico et al. 2007; Swope et al. 2014; Herbert et al. 2019). However, how cells might shape a tissue-restricted DNA-damage response is poorly understood.

Melanocytes, specialized pigment-producing cells in the skin, play a key role in photoprotection, but can also give rise to melanoma, the most lethal form of skin cancer, because of mutations that promote proliferation and suppress senescence (Shain and Bastian 2016). Much of melanocyte and melanoma biology is coordinated by the Microphthalmia-associated Transcription Factor (MITF) (Goding and Arnheiter 2019). MITF, a lineage survival oncogene (Garraway and Sellers 2006), controls cell survival (McGill et al. 2002)and autophagy (Ploper et al. 2015; Moller et al. 2019), promotes proliferation and differentiation (Widlund et al. 2002; Carreira et al. 2005; Loercher et al. 2005), but also suppresses invasion/migration (Carreira et al. 2006) and senescence (Giuliano et al. 2010; Ohanna et al. 2011). MITF also has a key role in regulating cell metabolism including mitochondrial biogenesis (Haq et al. 2013; Vazquez et al. 2013), the TCA cycle (Louphrasitthiphol et al. 2019) and fatty acid metabolism (Vivas-Garcia et al. 2020). Significantly, MITF can promote genome integrity by transcriptionally activating DNA damage repair genes, and depletion of MITF can trigger DNA damage (Beuret et al. 2011; Strub et al. 2011).

MITF is not a common target for melanoma-associated mutations. The notable exception, however, is a germline E318K mutation that is associated with increased melanoma and renal cell carcinoma predisposition in humans (Bertolotto et al. 2011; Yokoyama et al. 2011), and disease progression in a mouse melanoma model (Bonet et al. 2017). The E318K mutation prevents efficient SUMOylation of MITF K316 (Bertolotto et al. 2011; Yokoyama et al. 2011), but although it has been suggested that SUMOylation might affect the ability of MITF to regulate transcription (Miller et al. 2005; Murakami and Arnheiter 2005; Bertolotto et al. 2011) and suppress senescence (Bonet et al. 2017), mechanistically how SUMOylation affects MITF function is poorly understood. Nor is it known the range of biological processes controlled by SUMO-MITF.

Here we reveal that after UV or Gamma-irradiation, MITF is phosphorylated by ATM/DNA-PK and dissociates from its transcription co-factors. Phosphorylated MITF is recruited to sites of DNA damage, interacts with the MRN complex, limits HR-mediated repair and triggers replication stress, therefore increasing genome instability. Further, we describe how the E318K mutation recapitulates the effects of MITF phosphorylation and propose a new mechanism for the observed increase in melanoma predisposition.

## Results

### MITF expression correlates with replication stress and genome instability

Previous analysis of a potential link between MITF levels or activity and Single Nucleotide Variation (SNV) load failed to detect any significant correlation (Herbert et al. 2019). Here we used a different approach based on the rheostat model for MITF function (Carreira et al. 2006) that describes how low MITF is associated with invasiveness and a stem cell-like phenotype, moderate activity is found in proliferative cells, and high MITF corresponds to a more differentiated phenotype. Examination of the TCGA melanoma cohort stratified into 5 bins corresponding to different MITF levels provided an intriguing insight into the accumulation of SNVs in relation to MITF expression. These data show that both high (MITF^High^) and low (MITF^Low^) levels of MITF correlate with an increased mutational landscape (Fig. 1A). Previous work has indicated that MITF can control expression of genes implicated in DNA damage repair and consequently depletion of MITF can induce DNA damage (Beuret et al. 2011; Strub et al. 2011). Using the set of replication, recombination, and DDR-associated MITF-target genes described by Strub *et al*. (2011) we confirmed their downregulation upon MITF-targeting siRNA (siMITF) transfection into 501mel cells (Supplemental Fig. S1A). However, the increase in SNV load in MITF^High^ tumours was unexpected. We asked if an association between a change in MITF activity and accumulation of genomic alterations could also be observed in a non-pathological model comparing human pluripotent stem cell (hPSC)-derived melanoblasts, which do not express a range of MITF target genes associated with differentiation, and melanocytes which do. In this model, melanoblasts and mature melanocytes originate from the differentiation of the same population of hPSCs (Chambers et al. 2009; Lee et al. 2010; Mica et al. 2013; Baggiolini et al. 2021). Targeted sequencing of each population showed that only melanocytes displayed a significant amount of copy number alterations (CNA) while melanoblasts were similar to the control (Fig. 1B).

**Figure 1.**
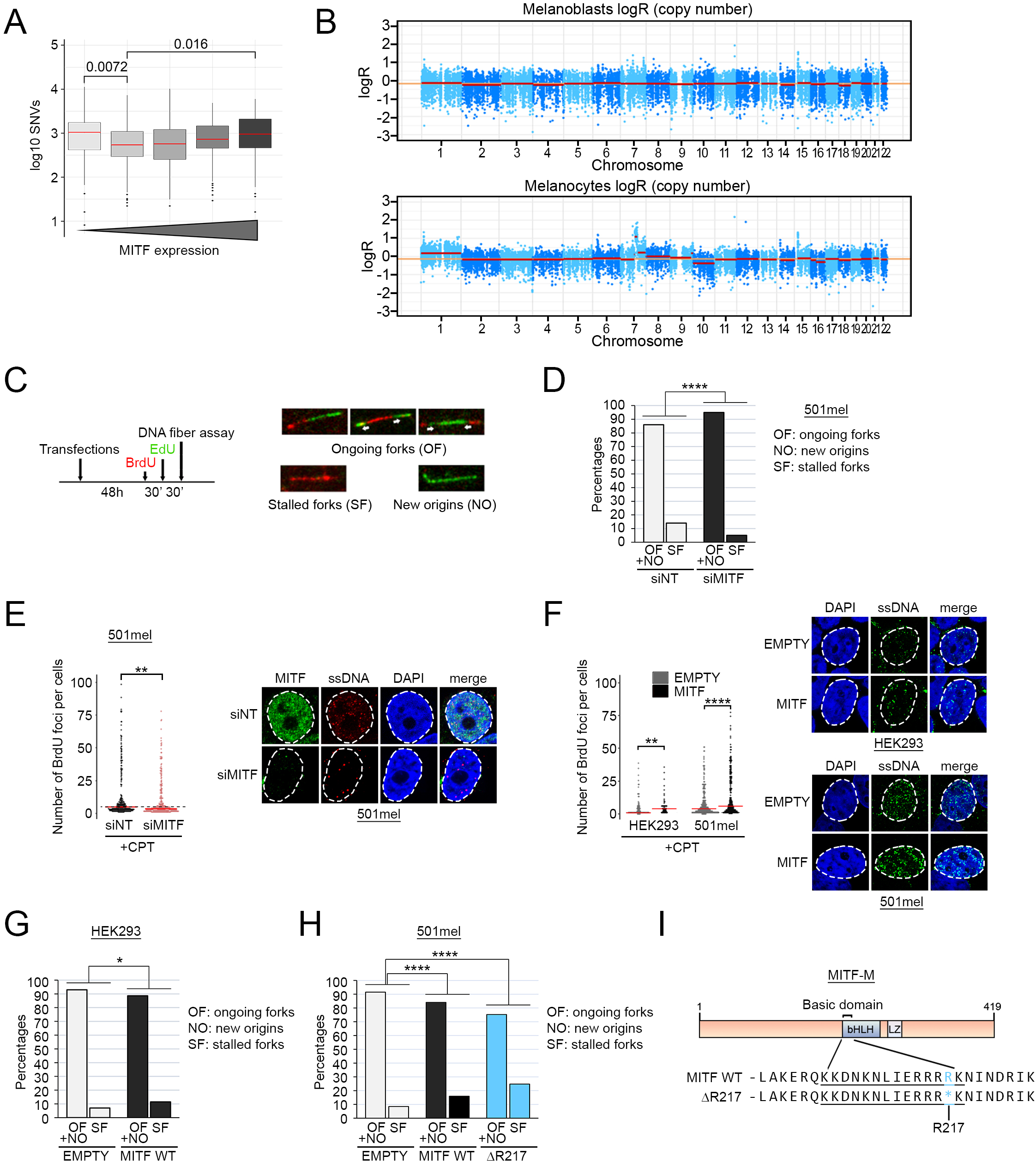
MITF associates with mutational burden and replication stress. (*A*) Boxplot showing the distribution of single nucleotide variants (SNVs) per sample, from 428 melanoma samples from ICGC, plotted after log-transformation computed as log10(0.1+SNV) in five bins by their MITF expression value. The *p-value* was computed using the paired Wilcoxon Rank Sum test. The medians are indicated in red. (*B*) Total copy number log-ratio (logR) of melanoblasts (top) and melanocytes (bottom). Orange horizontal line indicates the inferred diploid state and red lines show copy number segments along each chromosome. Melanoblasts displayed a flat profile indicating no copy number alterations whereas melanocytes showed copy number alterations on chromosomes 1,7 and 10. (*C* - *Left*) Timeline of the DNA fibre experiment. Cells were transfected for 48h before being treated sequentially 30 min with BrdU and 30min with EdU, before DNA extraction. (*C* - *Right*) Examples of patterns used to quantify the percentage of stalled forks, ongoing forks, and new origins in the DNA fibre assay. BrdU-containing fibres are stained in red, EdU-containing fibres are stained in green. (*D*) Graph expressing the percentage of ongoing forks (OF)/new origins (NO) and stalled replication forks (SF) in 501mel cells transfected with siMITF (black) or the non-targeting siRNA (grey). The *p-value* was determined using Fisher’s exact test. *p(siMITF)*= 9.453417e-06. (*E*) Immunofluorescence analysis of the accumulation of ssDNA in 501mel cells transfected with siMITF (grey) of the non-targeting siRNA (black). Cells incorporated BrdU for 24 h and were treated with 10 µM camptothecin (CPT) for 1 h. ssDNA foci were detected using an anti-BrdU antibody. *Left*: Beeswarm plots representing the quantification of the number of BrdU foci. The *p-value* was computed using the Wilcoxon Rank Sum test. *p(siMITF)*= 1.442e-06. The medians are indicated in red. *Right:* Representative images of ssDNA staining. (*F*) Immunofluorescence analysis of the accumulation of ssDNA after CPT treatment in HEK293 and 501mel transfected with HA-MITF or the corresponding empty vector. All cells had incorporated BrdU for 24 h and were treated with 10 µM camptothecin (CPT) for 1 h. ssDNA foci were made visible using an anti-BrdU antibody. *Left:* Beeswarm plots representing the quantification of the number of BrdU foci. The *p-values* were computed using the Wilcoxon Rank Sum test. *p(HEK293)*=0.002372; *p(501mel)*=6.014e-06. The medians are indicated in red. *Right*: Representative images of ssDNA staining in HEK293 *(top right panel)* and 501mel *(bottom right panel)*. (*G*) Graph expressing the percentage of ongoing forks (OF)/new origins (NO) and stalled replication forks (SF) in HEK293 cells overexpressing MITF WT (black) or the corresponding empty vector (grey). *P-values* were determined using Fisher’s exact test. *p(MITF WT)*= 0.0109008. (*H*) Graph expressing the percentage of ongoing forks (OF)/new origins (NO) and stalled replication forks (SF) in 501mel cells overexpressing MITF WT (black), ΔR217 (blue) or the corresponding empty vector (grey). *P-values* were determined using Fisher’s exact test. *p(MITF WT)*= 2.461506e-06; *p(del217)*= 1.960337e-19. (*I*) Diagram depicting the position of the basic domain of MITF and the ΔR217 deletion.

Since MITF can promote both cellular proliferation and differentiation (Widlund et al. 2002; Carreira et al. 2005; Loercher et al. 2005; Carreira et al. 2006), we asked if higher levels of MITF protein could trigger replication stress that might explain the genome instability we observed in melanocytes and melanoma. We performed a series of DNA fibre experiments in MITF-expressing 501mel melanoma cells. In this assay, cells are exposed first to BrdU for 30 minutes then EdU for the same duration to allow the two thymidine analogues to integrate into replicating DNA (Fig. 1C). After the second pulse, DNA is spread onto microscope slides, fixed and each analogue revealed using Click-It chemistry for EdU (green) and a specific antibody for BrdU (red). As a result, replicating DNA tracks can be visualised and discriminated into ongoing forks containing both BrdU and EdU, stalled forks that did not incorporate EdU (red only) and new origins that incorporated only EdU (green only). We first performed the DNA fibre assay in 501mel cells transfected with an MITF-targeting or a control siRNA (siCTL) (Fig. 1B and Supplemental Fig. S1B). Qualitative analysis showed a marked reduction of the percentage of stalled forks, from 15% with the control non-targeting siRNA to 5% with siMITF. This difference did not arise as a consequence of altered replication fork speed since the distribution of EdU track lengths in each sample did not reveal any difference (Supplemental Fig. S1C). In an orthogonal approach, we measured the accumulation of single-strand DNA (ssDNA) as a marker of replication stress after exposure to the topoisomerase poison camptothecin (CPT). siMITF or siCTL 501mel cells were allowed to incorporate BrdU for 24 h, then exposed to CPT and any exposed BrdU, reflecting ssDNA, was detected using a specific antibody under non-denaturing conditions. After quantification, the number of BrdU foci was lower in MITF-depleted cells, indicating a lower level of replication stress (Fig. 1E). We next transfected a HA-tagged MITF or a corresponding empty expression vector into the MITF-negative non-melanoma HEK293 cell line or MITF-positive 501mel melanoma cells and measured the amount of CPT-induced ssDNA and the DNA replication tracks. As before, MITF expression correlated with an increased amount of ssDNA in both HEK293 and 501mel cells (Fig. 1F) and a higher percentage of stalled replication forks (Fig. 1G and 1H). Taken together, the DNA fibre assay and the ssDNA detection experiments indicate that MITF expression correlates with the level of replication stress in melanoma cells.

### MITF-induced genome instability is independent of its transcriptional activity

Because it is unclear whether the difference in the percentage of stalled forks arose owing to the transcriptional activity of MITF, we generated a non-DNA binding MITF mutant by deletion of the arginine 217 (ΔR217) in the basic domain as described (Hemesath et al. 1994) (Fig. 1I). We then transfected 501mel cells with HA-MITF ΔR217 and performed the DNA fibre assay (Fig. 1H, blue bars). The results revealed 9% of stalled replication forks in the control cells, 16% in the MITF-transfected cells, and 25% with the ΔR217 mutant, indicating that replication fork stalling caused by MITF expression is independent of its ability to bind DNA and consequently its capacity to regulate transcription. Moreover, MITF expression was associated with globally decreased fork speed as shown by a shift of the peak of EdU track lengths to the left (Supplemental Fig. S1D), with an even greater shift being observed with MITF ΔR217 (Supplemental Fig. S1E). These results support the hypothesis that the effect of MITF on the replication machinery is non-transcriptional.

### MITF sensitizes cells to DNA damage

Knowing that MITF expression is associated with replication stress, we hypothesized that MITF-expressing cells would be more sensitive to DNA damage. We exposed MITF-negative HEK293 cells transfected or not with HA-MITF to UV, CPT, or the DNA alkylating agent cisplatin (CisPt). All three stresses have a deleterious effect on DNA replication that can lead to formation of DSB. Quantification of γH2AX, a marker of DNA damage, showed that expression of MITF increased γH2AX in response to UV, after exposure to CPT and to a lesser extent CisPt (Fig. 2A). We next performed a time course of γH2AX activation after UV in HEK293 cells transfected with HA-tagged MITF or a HA-only vector. We observed that more MITF-expressing cells displayed an elevated γH2AX signal 1 h after UV exposure, and that while by 24 h post-UV in the control cells the γH2AX signal was close to that in unirradiated cells, in the MITF-expressing population the γH2AX signal remained high (Fig. 2B). The experiment was repeated with the transcriptionally inactive MITF ΔR217 mutant and γH2AX detected over 4 h after UV irradiation (Fig 2c). The results revealed that the ΔR217 was similarly able to stimulate the activation of γH2AX in comparison to empty vector, even though its expression was slightly reduced compared to MITF WT by western blot (Fig. 2D). We concluded that direct DNA binding was not necessary for the enhanced induction of γH2AX observed using MITF WT following UV.

**Figure 2.**
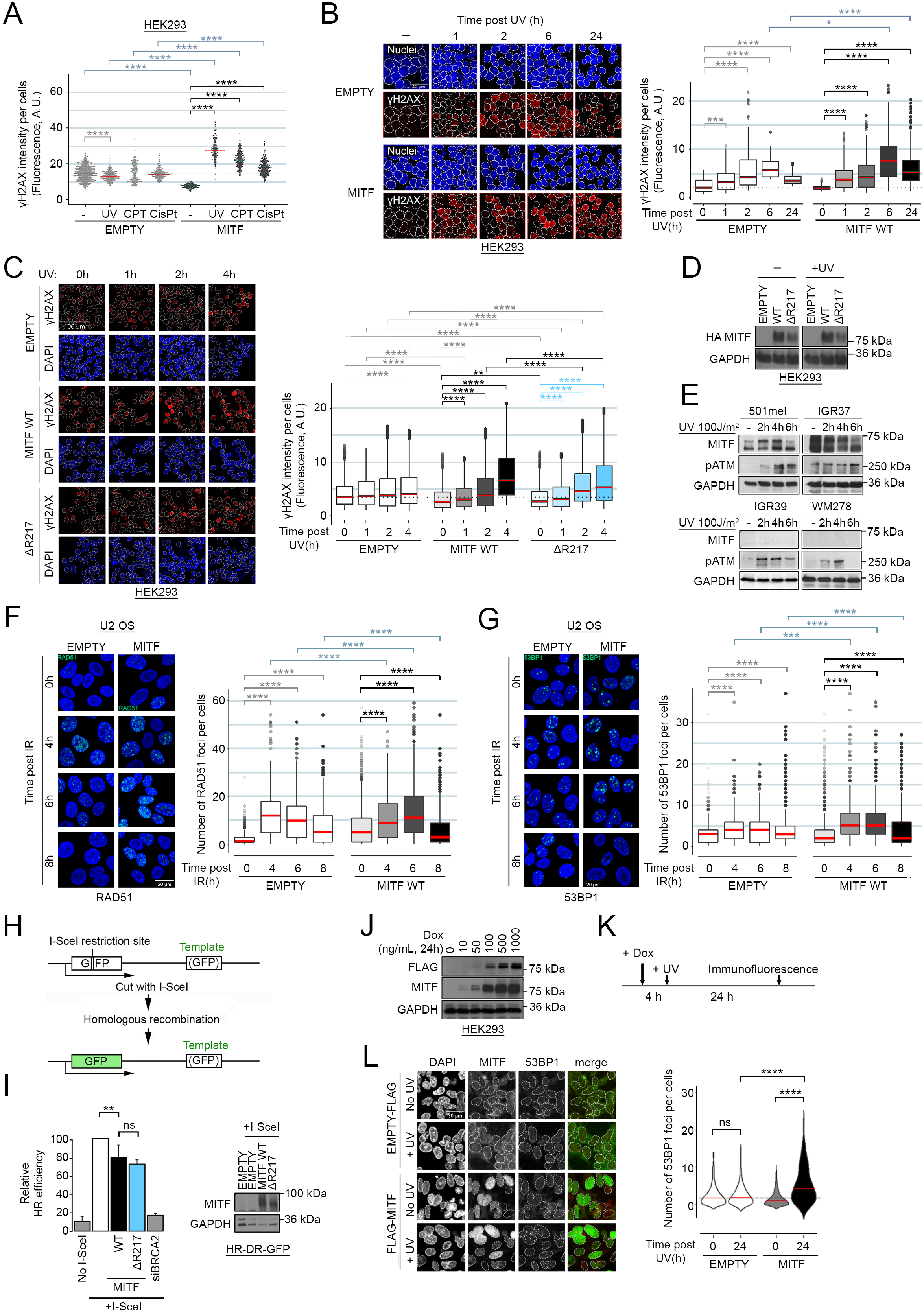
MITF positive cells are sensitive to different types of DNA damage. (*A*) Immunofluorescence analysis of γH2AX activation in HEK293 cells transfected with HA-MITF (black) or the corresponding empty vector (grey). Cells were exposed to 24 J/m^2^ UV, or treated for 2 h with 10 µM camptothecin (CPT) or 300 g/mL cisplatin (CisPt). Beeswarm plots representing the distribution of γH2AX intensities per cells. The *p-values* were computed using the Wilcoxon Rank Sum test. ****: *p<0.0001*. The medians are indicated in red. (*B*) Immunofluorescence analysis of γH2AX activation over time after UV irradiation (24 J/m^2^). HEK293 cells were transfected with HA-MITF (black) or the corresponding empty vector (grey). *Left*: representative images. *Right*: Boxplots representing the distribution of γH2AX intensities per cells. The *p-values* were computed using the Wilcoxon Rank Sum test. ***: *p<0.001;* ****: *p<0.0001*. The medians are indicated in red. (*C*) Immunofluorescence analysis of γH2AX activation over time after UV irradiation (24 J/m^2^). HEK293 cells were transfected with HA-MITF (black), the ΔR217 mutant (blue) or the corresponding empty vector (grey). *Left*: representative images. *Right*: Boxplots representing the distribution of γH2AX intensities per cells. The *p-values* were computed using the Wilcoxon Rank Sum test. **: *p<0.01;* ***: *p<0.001;* ****: *p<0.0001*. The medians are indicated in red. (*D*) Western blot of HEK293 cells transfected with HA-tagged MITF WT, the ΔR217 mutant or the control vector and exposed to UV (100 J/m^2^). GAPDH was used a loading control. (*E*) Western blot of melanoma cell lines expose to UV (100 J/m^2^)and harvested at the indicated times. Phospho-ATM was used as DDR activation control, GAPDH was used a loading control. (*F*) Immunofluorescence analysis of the formation of RAD51 foci after IR (X-rays, 2 Gy) in U2-OS cells transfected with HA-MITF or the corresponding empty vector. Cells were fixed and processed at the indicated times after IR. *Left*: representative images. *Right*: Boxplots representing the distribution of the number of RAD51 foci per cell. The *p-values* were computed using the Wilcoxon Rank Sum test. ****: *p<0.0001*. The medians are indicated in red. (*G*) Immunofluorescence analysis of the formation of 53BP1 foci after IR (X-rays, 2 Gy) in U2-OS cells transfected with HA-MITF or the corresponding empty vector. Cells were fixed and processed at the indicated times after IR. *Left*: representative images. *Right*: Boxplots representing the distribution of the number of 53BP1 foci per cell. The *p-values* were computed using the Wilcoxon Rank Sum test. ***: *p<0.001;* ****: *p<0.0001*. The medians are indicated in red. (*H*) Cartoon depicting the principle of the U2-OS-DR-GFP reporter system. (*I* - *left*) Graph showing the relative efficiency of homologous recombination using the U2-OS-DR-GFP reporter system. Data represent the mean (±SEM) from three independent experiments and are normalised against the I-SceI only samples. The *p-values* were computed using the Wilcoxon Rank Sum test. *ns*: non-significant; **: *p<0.01*. (*I* - *right*): Western blot of U2-OS-DR-GFP reporter cells transfected with MITF WT, the ΔR217 mutant or the control vector. GAPDH was used a loading control. (*J*) Western blot of FLAG-MITF induction with doxycycline. GAPDH was used a loading control. (*K*) Timeline of the experiment. Cells were induced with doxycycline for 4 h before being exposed to UV (24 J/m^2^). Immunofluorescence analysis was performed after 24 h. (*L*) Immunofluorescence analysis of the persistence of 53BP1 foci (red) in FLAG-MITF expressing cells (green). Violin plot representing the distribution of the number of 53BP1 foci per cells in inducible HEK293 cells expressing FLAG-MITF (grey) or the corresponding empty-FLAG (white) 24 h after being exposed to UV. The *p-values* were computed using the Wilcoxon Rank Sum test. *ns*: non-significant; ****: *p<0.0001*. The medians are indicated in red.

These data suggest that MITF levels affect the kinetics of the DDR. We therefore selected four melanoma cell lines, two MITF^High^ (501mel and IGR37) and two MITF^Low^ (WM278 and IGR39) and evaluated by western blot the dynamics of UV-induced ATM Ser1981 phosphorylation (pATM), a marker of active ATM (Fig. 2E). All cell lines displayed an activation of pATM within 2 h after UV exposure except for IGR37 where the kinase is already activated in steady state conditions. After the initial activation, pATM levels returned close to baseline 6 h after irradiation only in the MITF^Low^ cell lines WM278 and IGR39 while remaining high in 501mel and IGR37. Although only a correlation, this result is consistent with MITF expression perturbing the normal course of UV-induced damage repair.

As ATM and γH2AX are well-characterized markers of DSB repair, we investigated the involvement of MITF in the two major DSB repair pathways: HRR and NHEJ. First, we examined the formation of RAD51 foci to explore the impact of MITF expression on HRR activation in U2-OS cells, a well-defined model for the study of the formation of DDR foci and a cell line negative for MITF. U2-OS cells transfected with HA-MITF, or a control vector, were exposed to X-rays (IR) and the numbers of RAD51 foci was determined over time (Fig. 2F). With the control vector, the number of RAD51 foci reached a peak 4 h after IR and then started to decrease. In the presence of HA-MITF, the peak of RAD51 foci was delayed and observed only after 6 h. Note that the expression of MITF itself also increases the basal number of RAD51 foci. By 8 h post-IR, both samples displayed a similar recovery. We then asked if the delay in RAD51 foci formation would be compensated by an increase in other DSB repair mechanism and assessed the dynamics of formation of 53BP1 foci, a well-known marker of NHEJ-mediated repair. After IR, the number of 53BP1 foci increased between 4 h and 6 h in the control cells and started to decrease at 8 h (Fig. 2G). In the presence of MITF, the timing of activation was similar, however the number of foci per cell was higher than with the empty vector at both 4 h and 6 h. These results suggest that MITF expression triggers a shift from HRR to NHEJ in response to IR, possibly reflecting a deficiency in HRR. To confirm this, we used the U2-OS-DR-GFP cell reporter system (Pierce et al. 1999). In this assay, engineered U2-OS cells contain a GFP cDNA containing a blunt-ended I-SceI digestion site and another fragment of GFP spanning the cleavage site. After enzymatic digestion with I-SceI, cells use the second GFP fragment as a template to restore a functional fluorescent GFP (Fig. 2H). We co-transfected the reporter cells with expression vectors for I-SceI and MITF or an empty vector and measured GFP expression (Fig. 2I). In the control cells GFP expression was markedly increased upon expression of I-Sce1. Co-expression of MITF WT moderately, but consistently reduced the efficiency of HRR by approximately 20% compared to I-SceI alone. As a control, depletion of BRCA2 decreased HRR efficiency to around 20%. Notably, the non-DNA binding MITF-ΔR217 mutant also reduced the efficiency of HRR, confirming that the effects of MITF on HRR are likely non-transcriptional.

To confirm these observations, we next used a HEK293 cell line stably expressing a FLAG-tagged MITF (or its empty FLAG counterpart) under a doxycycline-inducible promoter. Using a range of doxycycline concentrations for 24 h, we determined that FLAG-MITF was induced at 10 ng/mL and its expression increased with increased doxycycline (Fig. 2J). We chose to work with 100 ng/mL doxycycline to induce an intermediate level of MITF consistent with that observed in MITF^High^ melanoma cell lines. For the persistent damage assay, we induced FLAG-MITF for 4 h prior to exposing cells to UV and then allowed cells to rest for 24 h while maintaining MITF induction with doxycycline. Cells were then analysed by immunofluorescence for persistent 53BP1 foci indicative of unrepaired DSB (Fig. 2K). 24 h after UV, no significant accumulation of 53BP1 foci was observed in the FLAG-only cells, while FLAG-MITF-expressing cells displayed a significant increase in persistent 53BP1 foci (Fig. 2L). This result confirmed that MITF expression decreases the efficiency of repair of UV-induced DNA damage leading to the accumulation of unrepaired damage.

### DNA damage remodels the MITF interactome

Our results so far established that MITF delays the resolution of UV-induced damage without the need to directly bind DNA. We therefore hypothesized that MITF may have a non-transcriptional function in the regulation of DNA repair by interacting with known DNA repair factors. To test this, a stable HEK293 cell line expressing a doxycycline-inducible BirA*-FLAG-MITF (Chauhan et al. 2022) was used in an unbiased search for MITF interactors using biotin-streptavidin-based purification coupled to mass spectrometry (BioID) (Lambert et al. 2015). BioID was performed 20 h after induction of MITF using doxycycline in the presence or absence of CPT. This identified 70 MITF-interacting proteins in the absence of CPT including the other members of the MiT family TFEB and TFE3 that can heterodimerise with MITF (Chauhan et al. 2022) (Fig. 3A and 3B).

**Figure 3.**
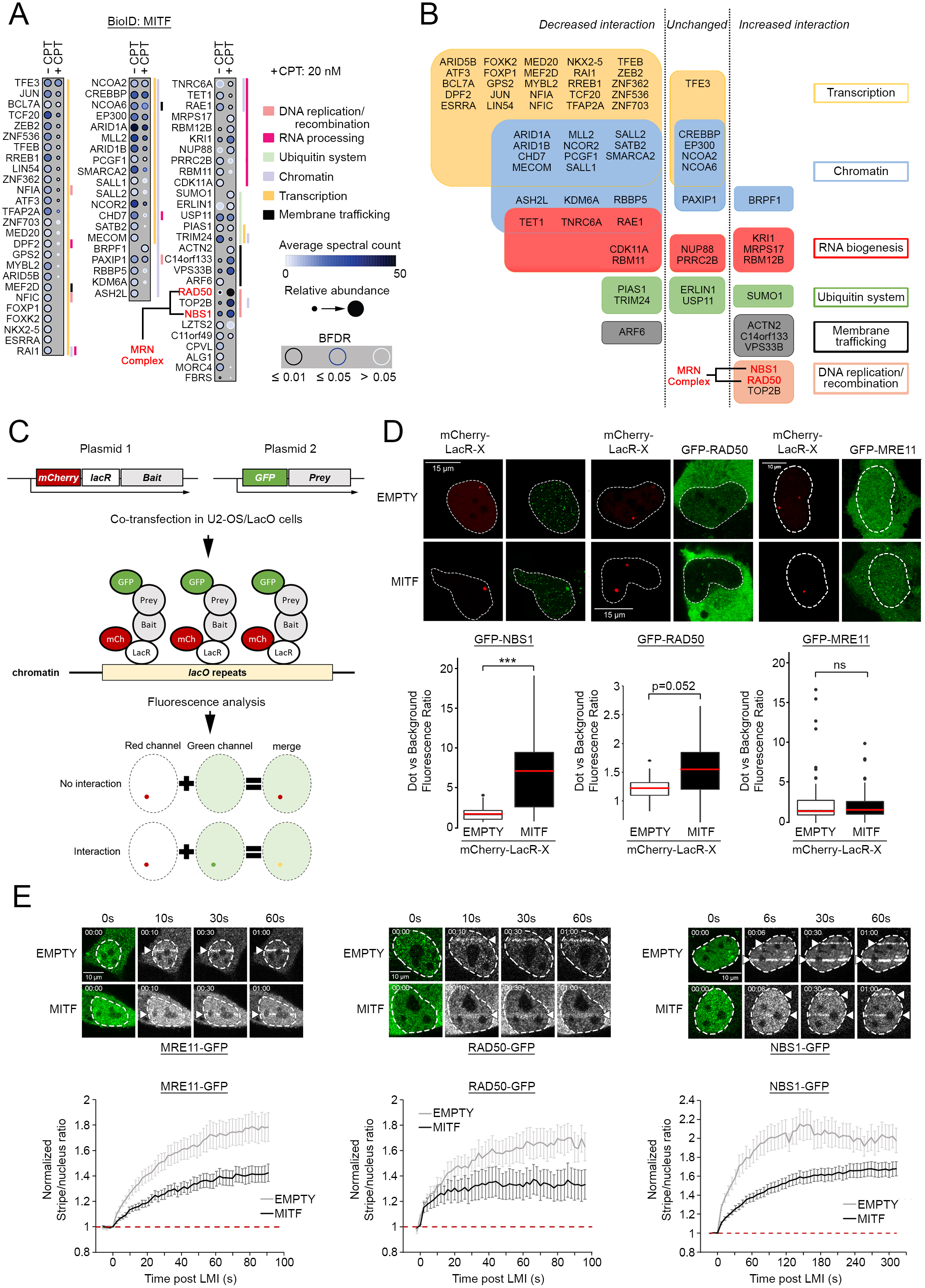
MITF interactome is remodelled by DNA damage. (*A*) Dot plot of BioID data showing the significant proximity partners of MITF in non-treated versus CPT-treated HEK293 cells. The colour of the dots represents the average spectral count. The size of the dots represents the relative abundance between conditions. The grey intensity of the line encircling the dots represents the Bayesian False Discovery Rate (BDFR) cut off. Association to either of the six categories: DNA replication and recombination, RNA processing, Ubiquitin system, Chromatin, Transcription and Membrane trafficking is depicted by the coloured lines to the right of each box. Components of the MRN complex are indicated in red. (*B*) Diagram representing the BioID results according to the effect of CPT on MITF interaction. Where possible, proteins from Fig. 3A were classified using the KEGG BRITE database (www.kegg.jp) and information from UNIPROT (www.uniprot.org) into six categories: transcription regulation; chromatin regulators; RNA processing; ubiquitin system; membrane trafficking; and DNA replication and recombination and segregated into columns highlighting (from left to right) a decreased, unchanged or increased interaction with MITF after CPT treatment. Components of the MRN complex are indicated in red. (*C*) Cartoon depicting the nuclear tethering assay. (*D* - *Top*) Representative images and quantification of the nuclear tethering assay showing interaction between MITF and NBS1, RAD50 or MRE11. The left panels show the localization of mCherry-LacR-NLS or mCherry-LacR-MITF dots in the nuclei of U2OS-LacO#13 cells, and the right panels show GFP-NBS1, GFP-RAD50 or GFP-MRE11 respectively. (*D* - *Bottom*) The quantification is expressed as the ratio between the GFP fluorescence measured inside the area delimited by the mCherry dot and in the rest of the nucleus. The *p-values* were computed using the Wilcoxon Rank Sum test. *ns*: non-significant; ***: *p<0.001*. The medians are indicated in red. (*E* - *top*) Still images of MRE11-GFP, RAD50-GFP and NBS1-GFP LMI. Nuclei are delimited using the pre-irradiation images. The positions of the irradiated lines are indicated with an arrow. (*E* - *bottom*) Quantification of MRE11-GFP, RAD50-GFP and NBS1-GFP recruitment in U2-OS cells after LMI when co-transfected with an empty vector or HA-MITF. The graphs represent the mean +/- SEM stripe/nucleus ratio over time. Values are normalized against the pre-LMI measurements. The baselines are indicated with red dotted lines.

The majority of MITF interaction partners are transcription factors and presumably represent DNA binding partners that might facilitate high affinity MITF binding to sequence elements recognised by both proteins. This set includes TFAP2A, a transcription factor that genetically interacts with MITF to control melanocyte development and differentiation and which genome wide co-occupies a subset of sites bound by MITF (Seberg et al. 2017; Kenny et al. 2022). The second most represented proteins are chromatin regulators, including the histone acetyltransferases CREBBP and p300, known coactivators for MITF (Sato et al. 1997; Price et al. 1998), but which also mediate MITF acetylation (Louphrasitthiphol et al. 2020). Also identified was SMARCA2 a member of the SWI/SNF complex that functions as a MITF co-factor (de la Serna et al. 2006; Saladi et al. 2013; Laurette et al. 2015), as well as several other potential co-factors not previously identified as MITF interactors. MITF also interacts significantly with RNA-binding and processing factors suggesting a new role in RNA biology beyond transcription. We confirmed the previously known interaction with the de-ubiquitinase USP11 and the E3-ubiquitin ligase TRIM24 (Laurette et al. 2015), as well as the E3 SUMO-protein ligase PIAS1 (Murakami and Arnheiter 2005). The BioID analysis also discovered lysosome-associated factors (VPS33B/C14orf133) in line with the association of some MITF family members with the lysosome (Martina and Puertollano 2013) and the role of MITF in autophagy and lysosome biogenesis (Moller et al. 2019). Also found, albeit less represented, were factors involved in cytokinesis ARF6, LZTS2 and RAE1, consistent with a role for MITF in spindle formation and chromosome segregation (Strub et al. 2011). More interestingly, given our results showing potential MITF regulation by DNA-PK, we revealed interactions between MITF and the DNA repair protein PAXIP1, which is also involved in transcription regulation, as well as RAD50 and NBS1. Together with MRE11, RAD50 and NBS1 are both members of the MRN complex that plays a major role replication fork restart as well as in the early steps of DSB repair and which is required for the activation of ATM (Uziel et al. 2003).

After treating cells with CPT, only 10 of the 70 interacting proteins detected were found equally in both control and CPT conditions including TFE3, CREBBP, EP300, NCOA2 and NCOA6. Remarkably, apart from TFE3 that can dimerize with MITF, all transcription factors and most chromatin regulators exhibited decreased interaction with MITF, suggesting that MITF transcriptional activity is attenuated. An exception was BRPF1 that regulates the acetylation of histone H3K23, that showed a significant association with MITF only after CPT treatment, but which has not been previously linked to the DDR. RNA processing factors KRI1 and MRPS17, associated with the ribosome biogenesis, are also overrepresented after CPT treatment. Notably we saw decreased interaction with TET1, a methyl cytosine dioxygenase implicated in DNA-demethylation, but which is associated with spliceosomes. We noted that SUMO1 is detected only after CPT treatment. Since SUMOylated peptides are difficult to detect using mass spectrometry, our interpretation is that we may be detecting SUMO1 that is interacting with, rather than modifying MITF.

### MITF interacts with components of the MRN complex

Of the MITF-interacting partners that were more abundant after damage those implicated in the DNA replication and DDR, TOP2B, RAD50 and NBS1, were especially interesting. TOP2B is a type II topoisomerase required to relax the topology of the DNA double helix prior to transcription and replication and is not inhibited by CPT. Given the potential for MITF to function in DNA replication and damage repair, we focused on its interaction with RAD50 and NBS1. Intriguingly, while NBS1, RAD50 and MRE11 function in a ternary complex, we never observed MRE11 as a significant MITF proximity partner (SAINTexpress FDR ≤ 1%).

To confirm the BioID results in an *in vivo* setting using an orthogonal approach, we examined the interaction between MITF and components of the MRN complex using the LacR/LacO nuclear tethering system (Luijsterburg et al. 2012). This live cell-based assay uses a U2-OS cell line containing an array of 256 repeats of the Lac operator (LacO) DNA sequence integrated into the genome, used to tether a bait fusion protein containing the Lac Repressor (LacR) tagged with mCherry to produce a single red fluorescent nuclear focus in G1 cells (Lambert et al. 2019). Cells are then transfected with the putative interactor protein tagged with GFP and co-localization of the GFP-fusion with the mCherry nuclear focus indicates interaction (Fig. 3C). Since MITF can form homodimers, by using mCherry-LacR-MITF as bait we first verified that we could detect an interaction using MITF-GFP as a positive control. The results (Supplemental Fig. S2A) showed a clear co-localization of tethered mCherry MITF with MITF-GFP consistent with MITF dimerization. No significant interaction was observed if we used LacR-mCherry-MITF with GFP only, or with both empty vectors. Using again LacR-mCherry-MITF as bait we next monitored the interaction with GFP-tagged NBS1, RAD50 or MRE11. Only NBS1-GFP was primarily nuclear and accumulated significantly at the MITF-rich nuclear mCherry dot with a median enrichment of 7-fold (Fig. 3D, left panel). No co-localization of NBS1-GFP was detected using an mCherry bait lacking MITF. RAD50-GFP, despite its predominantly cytoplasmic expression, also displayed interaction (Fig. 3D, middle panel). On the other hand, MRE11-GFP did not accumulate at the LacO array irrespective of whether we used the control vector or mCherry-LacR-MITF (Fig. 3D, right panel). This last observation supports the results from the BioID experiment where MRE11 was not detected as a significant MITF proximity partner. We also controlled the interaction between each member of the MRN complex (Supplemental Fig. S2B). As expected, MRE11-GFP recruitment was enriched in the presence of mCherry-LacR-RAD50 or NBS1, but the interaction between NBS1-GFP and mCherry-LacR-RAD50 was not significant. These observations reflect the dynamics of interactions inside the MRN complex as described previously (Lafrance-Vanasse et al. 2015).

Under steady state conditions, MRE11 and RAD50 form a cytoplasmic complex with NBS1 that is required to bring the complex into the nucleus (Tauchi et al. 2002). While the possibility of a complex between RAD50 and NBS1 lacking MRE11 has only been described *in vitro* (van der Linden et al. 2009), the physical interaction between MITF and RAD50/NBS1 but not MRE11 prompted us to ask if it was of functional significance. Knowing that MITF limits HRR activation and that the MRN complex is implicated in DSB repair pathway choice by promoting HRR and inhibiting NHEJ, we asked if interacting with MRN MITF could affect its recruitment to DSB and thereby impair HRR. We used laser microirradiation (LMI) with a near-infrared (NIR) laser emitting at 750 nm, to induce damage in a strictly defined area of the nucleus and performed a series of NIR-LMI experiments coupled with video microscopy in U2-OS cells that lack endogenous MITF to analyse the dynamics of recruitment to DNA damage of the different components of the complex (Supplemental Fig. S2C). We then compared the relative recruitment of each factor. All members of the MRN complex are recruited rapidly in a few seconds but their recruitment was substantially impaired by co-expression of MITF WT (Fig. 3E and Supplemental Videos 1, 2 and 3).

### ATM- and DNAPK-dependent MITF recruitment to DNA damage

Given the non-transcriptional effect of MITF on γH2AX activation, and MITF’s interaction with the MRN complex, we next asked whether MITF could be directly involved in the DDR pathway using 501mel cells stably expressing HA-MITF at endogenous levels (Louphrasitthiphol et al. 2020). We performed LMI using a UV-emitting laser and detected the localisation of MITF by immunofluorescence. Twenty min after UV-LMI, MITF colocalised with γH2AX (Fig. 4A). To measure more precisely the dynamics of MITF recruitment, we repeated this experiment and irradiated adjacent regions every 10 s for 15 min before staining. We observed that GFP-MITF accumulated at the damage sites by 100 s after UV (Supplemental Fig. S3A). We repeated the experiments with NIR-LMI and confirmed that, like with UV, HA-MITF accumulated at the sites of damage (Fig. 4B). We then assessed the role of key DDR factors in MITF recruitment. We generated a stable 501mel cell line expressing GFP-MITF and performed live confocal microscopy after NIR-LMI. This revealed that recruitment of MITF to DNA damage was prevented by the DNA-PK inhibitor NU7441 (Fig. 4C, Supplemental Video 4), but not by the ATM inhibitor KU55933 nor ATR inhibitors (VE821 and VE822) (Supplemental Fig. S3B, Supplemental Video 5). Inhibition of poly-ADP ribose polymerase (PARP) using Olaparib also prevented MITF recruitment to DNA damage (Fig. 4C, Supplemental Video 6).

**Figure 4.**
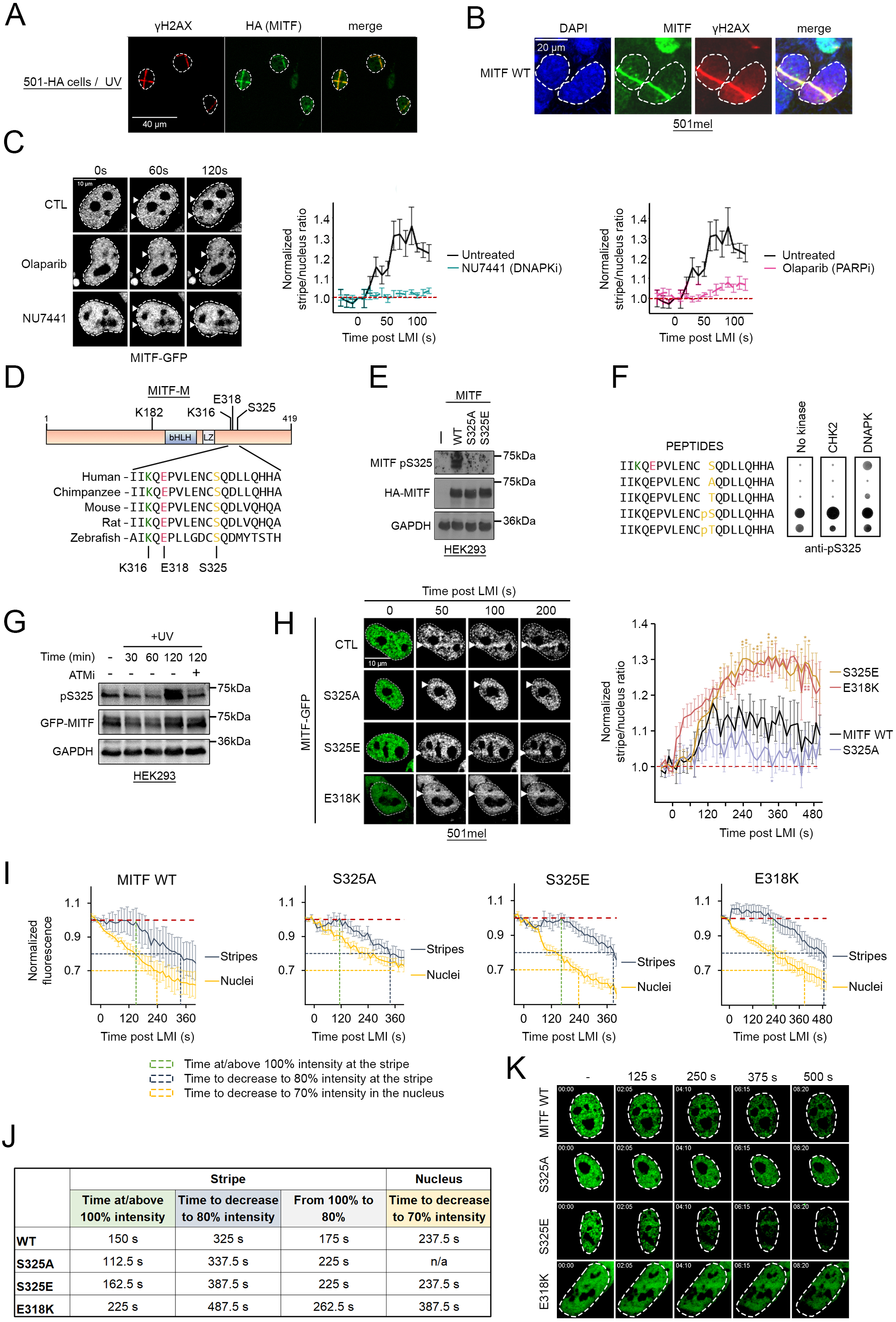
MITF recruitment to DNA damage sites. (*A*) Immunofluorescence of stable 501HA-MITF cells after UV-LMI. The irradiated area is identified using an anti-γH2AX antibody (red channel). MITF is detected with an anti-HA antibody (green channel). (*B*) Immunofluorescence of stable 501HA-MITF cells after NIR-LMI. The irradiated area is identified using an anti-γH2AX antibody (red channel). MITF is detected with an anti-HA antibody (green channel). DAPI was used to stain the nuclei. (*C*) Still images and quantification of live video-microscopy showing recruitment of MITF in 501mel cells stably expressing GFP-MITF after NIR-LMI. Cells were treated with DNA-PK (NU7441 – 1 µM) or PARP (Olaparib – 10 µM) inhibitors for 24 h before irradiation. The graphs represent the mean +/- SEM stripe/nucleus ratio over time. Values are normalized against the pre-LMI measurements. For clarity, each individual graph represents the same control curve (black) against only one type of inhibitor (DNAPKi in turquoise and PARPi in pink). The baselines are indicated with red dotted lines. (*D - top*) Diagram depicting the position of the bHLH-LZ domain of MITF and the position of the SUMOylation sites K182 and K316 (green), the phosphorylation site S325 (red) and the familial mutation E318K (pink). (*D - bottom*) Sequence of the 314-333 peptide showing residues K316, E318 and S325 are conserved from Human to Zebrafish. (*E*) Western blot of HEK293 cells transiently transfected with various mutants of HA-MITF or the corresponding empty vector. Phosphorylation of MITF on Serine 325 was detected by a phospho-specific antibody. GAPDH was used a loading control. (*F*) *In vitro* kinase assay performed on a peptide array. Peptide-bound anti-pS325 antibodies were detected using chemiluminescence. The size and intensity of the dots are proportional to the amount of bound antibodies. The kinases used are indicated above. (*G*) Western blot of HEK293 cells transiently transfected with HA-MITF. Cells were UV-irradiated and harvested at different time points to measure phosphorylation of S325. The ATM inhibitors KU55933 was used to confirm the role of the kinase. GAPDH was used a loading control. (*H - Left*) Still images of GFP-MITF recruitment in stable 501GFP-MITF WT, S325 and E318 mutants cell lines after LMI. Nuclei are delimited using the pre-irradiation images. The positions of the irradiated lines are indicated with an arrow. (*H - Right*) Quantification. The graph represents the mean +/- SEM stripe/nucleus ratio over time. Values are normalized against the pre-LMI measurements. The baseline is indicated with a red dotted line. (*I*) Details of GFP-MITF behaviour in stable 501GFP-MITF WT, S325 and E318 mutants cell lines after LMI. Quantification of GFP intensities at (stripes) and away from the LMI sites (nuclei). The graph represents the mean +/- SEM GFP fluorescence over time. Values are normalized against the pre-LMI measurements. The dotted lines represent the key values as described in (*J*). The baselines are indicated with red dotted lines. (*J*) Key values extracted from the quantification in (*I*). *Stripe – time at/above 100% intensity* represents the time GFP fluorescent remained equal or above 1 after LMI; *stripe – time to decrease to 80% intensity* represents the time required to see the GFP intensity at the stripe drop to 80% of the pre-LMI value from the time of irradiation; *stripe – from 100% to 80%* represents the time required to see the GFP intensity at the stripe drop to 80% of the pre-LMI value after it started to decrease; *nucleus – time to decrease to 70% intensity* represents the time required to see the GFP intensity away from the stripe drop to 70% of the pre-LMI value. (*K*) Still images of GFP-MITF behaviour in stable 501GFP-MITF wild type, S325 and E318 mutants cell lines after LMI as quantified in **i**. Nuclei are delimited using the pre-irradiation images.

Considering the effect of the DNA-PK inhibitor on MITF recruitment, we asked whether MITF might be a direct target of the kinases implicated in DDR. By examining evolutionarily conserved sequences within MITF we identified, C-terminal to the MITF Leucine Zipper, a highly conserved SQ motif (Fig. 4D) characteristic of targets of the ATM and DNA-PK kinases (Kim et al. 1999). Initial attempts to identify phosphorylation at this residue using a mass spectrometry approach were unsuccessful since although post-translational modifications at many other sites were identified, we were unable to generate peptide coverage of the region containing S325. As an alternative strategy, we generated a phosphorylation-specific polyclonal antibody against phospho-S325 and validated its specificity using a peptide array (Supplemental Fig. S3C) spanning mouse or human MITF amino acids 314-333 that contained either the WT sequence, S325 variants or, as an additional control, acetyl-K316. The results (Supplemental Fig. S3D) confirmed that the antibody was highly specific, recognising pS325 in both the mouse or human sequence, but neither the non-phosphorylated version, S325A or S325T mutants. A pT325 variant was recognised less well than pS325, and acetylation at K316 did not affect antibody recognition. To confirm the specificity of the anti-pS325 antibody in a cellular context, we transfected HEK293 cells that do not express endogenous MITF with plasmids expressing HA-tagged MITF WT or S325A (null) and S325E (phosphomimetic) phosphorylation site mutants. Following transfection cells were analysed by western blot using the pS325 antibody. The results (Fig. 4E) revealed that MITF WT, but not the S325 mutants were recognised by the anti-pS325 antibody confirming both the specificity of the antibody for pS325 and that S325 is phosphorylated in cells.

We next investigated the ability of DDR kinases to phosphorylate MITF S325 using two different approaches. First, we used commercially available purified CHK2 and DNA-PK in a non-radioactive *in vitro* kinase assay on a peptide array. In this assay, only the DNA-PK substantially increased the signal on the non-phosphorylated S325 residue and weakly on T325 (Fig. 4F). Because commercially available purified ATM was unavailable, we next used a specific ATM inhibitor KU-55933 to treat HEK293 cells transfected with HA-tagged MITF prior to UV irradiation and examined the extent of MITF S325 phosphorylation over time by western blot (Fig. 4G). The results confirmed that S325 phosphorylation can be detected in cells and increases after UV irradiation. The western blot also revealed that the ATM inhibitor was able to reduce the levels of S325 phosphorylation 2 h after UV. These results suggest that both ATM and DNA-PK can target MITF S325 for phosphorylation.

We asked whether phosphorylation of S325 could affect MITF recruitment to DNA damage. We generated a series of stable 501mel cell lines expressing GFP-tagged MITF WT and mutants and performed live cell NIR-LMI. We then quantified the dynamics of recruitment of GFP-MITF WT, S325A and S325E mutants (Fig. 4H, Supplemental Video 7). GFP-MITF WT accumulated slowly at the damage sites and becomes detectable after 30 s, reaching a maximum intensity between 120 s and 240 s. By comparison, the recruitment of the MRN complex, early factors in DSB repair, happens in as little as 5 s as shown in Fig. 3E. Compared to MITF WT, recruitment of the S325A mutant was significantly reduced. By contrast, the S325E mutant exhibited substantially enhanced recruitment, reaching a maximum at 360 s, though already at 240 s its presence at the damage stripe was well above that of the WT protein. These data suggest that phosphorylation at S325 represents a key regulator of MITF recruitment to DNA damage sites. Since MITF recruitment is dependent on DNA-PK activity that may be one of the kinases responsible for MITF S325 phosphorylation, we postulated that the inhibitor would have a reduced effect on the S325E mutant compared to MITF WT. We repeated the NIR-LMI experiment on 501mel GFP-MITF S325E cells pre-treated with the DNA-PK inhibitor or Olaparib as a positive control. The DNA-PK inhibitor NU7441 was still effective at reducing MITF S325E recruitment but less potent compared to MITF WT, supporting the role of the DNA-PK in MITF phosphorylation (Supplemental Fig. S3E, Supplemental Video 8). As expected, the PARP inhibitor Olaparib abolished recruitment.

Interestingly, residue S325 is close to the germline mutation site E318K known to increase melanoma risk (Bertolotto et al. 2011; Yokoyama et al. 2011) and prevent SUMOylation on K316 to affect the transcriptional program of MITF. Given the proximity of S325 with K316 we generated a 501mel GFP-MITF E318K cell line and measured the recruitment of MITF E318K to the NIR-LMI generated damage (Fig. 4H). Strikingly, MITF E318K was rapidly recruited, and the quantification showed a similar dynamic to what was observed with MITF S325E, with both mutations exhibiting a significantly increased recruitment compared to MITF WT, suggesting that SUMOylation on K316 is also a regulator of MITF function in the DDR.

In closely examining the dynamics of MITF recruitment we observed that the increased stripe/nucleus ratio was mainly due to a rapid decrease in the amount of GFP-MITF away from the stripe immediately after LMI. To confirm this, we measured separately the intensity of GFP-MITF at the damage sites (Fig. 4I, blue curves) and the intensity away from damage (Fig. 4I, yellow curves). The graph shows that MITF stably stayed at the stripe for approximately 150 s before diminishing (green dotted line) while its nuclear intensity decreased immediately after irradiation (Fig. 4I, left panel). To compare the behaviour of all the mutants, we marked the time required to reach a 20% decrease in fluorescence at the stripe (dotted blue lines) and a 30% decrease in the nucleus (dotted yellow lines). MITF WT stripe intensity decreased to 80% in 325 s and its nuclear intensity decreased to 70% in 237.5 s (Fig. 4I and 4J). In comparison, the MITF S325E mutant is more stable at the stripe (162.5 s at 100% and 387.5 s to 80%) and MITF S325A is more stable in the nucleus (fluorescence value remained over 70% during the course of the experiment) showing that S325 phosphorylation helps stabilize MITF at the stripe while promoting a decrease in the nucleus. These observations explained the previously noted differences in the ‘recruitment’ dynamics (Fig. 4H). On the other hand, the SUMOylation mutant E318K is the only mutant showing increased recruitment and stability at the stripe up to 225 s post LMI. At the same time, the decrease of E318K fluorescence in the nucleus is slower than the WT (30% decrease in 387.5 s). We also analysed the LMI data of MITF in the presence of the DNA-PK inhibitor NU7441. Similar to the S325A mutant, MITF was more stable in the nucleus in DNA-PK inhibitor-treated cells (20% decrease in 100 s in NU7441-treated against 50 s in control) but is rapidly lost at the stripe, with the two curves overlapping over the course of the experiment (Supplemental Fig. S3F). These results suggest that the main contributor to the apparent accumulation of MITF after DNA damage is the stability of the protein. Using the protein synthesis inhibitor cycloheximide (CHX) to arrest de novo MITF synthesis, we measured the half-life of endogenous MITF after treatment with inhibitors of ATM and DNA-PK, the two kinases able to phosphorylate MITF S325 and observed that both inhibitors reduced MITF protein stability (Supplemental Fig. S3G). Taken together, these results suggest that phosphorylation of MITF at sites of DNA damage contributes to its stabilization, and that MITF is less stable in absence of DNA-damage-induced phosphorylation.

### MITF S325E mutant phenocopies the MITF E318K

Since both S325 phosphorylation and K316 SUMOylation regulate the role of MITF in HRR and replication stress, it was possible that one modification may regulate the other. Examination of the amino acid sequence of human MITF in the vicinity of S325 revealed that the sequence I-K-Q-E-P-V-L-E-N-C-S-Q-D-D resembles a Negatively charged amino acid–Dependent SUMO-conjugation Motif (NDSM) ψ-K-x-E-x-x-x-E-x-x-S-x-D-D raising the possibility that ATM-induced S325 phosphorylation could regulate MITF K316 SUMOylation (Hietakangas et al. 2006; Yang et al. 2006). This is important given the role of the SUMOylation-defective E318K melanoma predisposition mutant identified previously (Bertolotto et al. 2011; Yokoyama et al. 2011). Using SUMO1-RFP or SUMO2-RFP fusion proteins co-expressed with HA-MITF, we confirmed that in HEK293, SUMO1 is more able to modify MITF as seen with a slow-migrating MITF band corresponding to SUMOylated MITF only when SUMO1-RFP was present (Supplemental Fig. S4A). Because MITF can be modified by either SUMO1 or SUMO2 (Miller et al. 2005; Murakami and Arnheiter 2005; Bertolotto et al. 2011; Yokoyama et al. 2011), we also used the deconjugation-defective mutant SUMO2-Q90P (Bekes et al. 2011; Garvin et al. 2013) to confirm that SUMO2 is able to target MITF but with lower efficiency. We generated a HA-MITF K182R expression vector which had no effect on the SUMO-MITF band confirming that SUMOylation must happen most favourably on K316 with WT HA-tagged MITF in HEK293 cells. We then generated double mutants of MITF to study the relationship between S325 phosphorylation and K316 SUMOylation. The introduction of either S325 mutation did not impact the level of MITF SUMOylation.

Previous work reported that K316 SUMOylation can affect MITF transcriptional activity on specific promoters, and that E318K mutant cells are more proliferative than their wildtype counterparts (Bertolotto et al. 2011). We therefore asked if the MITF S325E mutant, which behaves like E318K in the context of DNA and replication fork progression, would also recapitulate the effect of E318K on transcription and proliferation. To evaluate this, we co-transfected MITF WT or mutants with a *MET*-promoter luciferase reporter, a well-characterised proliferation-related target for MITF (McGill et al. 2006). After adjusting for the expression levels of MITF WT and mutants, we confirmed that the *MET* promoter is activated by MITF WT (Supplemental Fig. S4B), as well as by the MITF S325A mutant that permits MITF SUMOylation at K316. By contrast, the SUMOylation-defective E318K mutant reduced reporter activation, in accordance with previous observations (Bertolotto et al. 2011), as did the phospho-mimetic S325E mutant that impairs K316 SUMOylation. Using a colony formation assay in which cells were plated at low density, expression of the SUMOylation-defective E318K mutant gave a growth advantage to HEK293 cells compared to cells expressing MITF WT, as previously described (Bertolotto et al. 2011). The S325E mutant also displayed a significant increase in colonies compared to MITF WT or the S325A mutant (Supplemental Fig. S4C). We performed the same experiment in the 501mel melanoma cells and obtained a similar result in which the S325E and E318K mutants exhibited an increased growth rate compared to MITF WT and the S325A mutant (Supplemental Fig. S4D). Overall, our data revealed that the MITF S325E phospho-mimetic mutant recapitulates much of the effect of the germline E318K SUMOylation-defective mutant on transcription and proliferation.

### MITF phosphorylation and SUMOylation regulate its role in Homologous recombination-mediated repair

Given the roles of S325 and K316/E318 in controlling MITF recruitment to DNA damage we next investigated the effect of their respective mutations on HRR. Examination by immunofluorescence of RAD51 foci after gamma irradiation in U2-OS cells transfected with HA-MITF, HA-MITF-E318K or the corresponding control plasmid at 4 h and 6 h post-irradiation (Fig. 5A) revealed that in the control cells the accumulation of RAD51 foci peaked at 4 h before decreasing at 6 h. By contrast, expression of either HA-MITF or MITF-E318K provoked a delay, with RAD51 foci reaching a maximum at 6 h. In the same experiment, the phospho-mimetic S325E also delayed the accumulation of RAD51 foci, with no increase observed at 4 h, while the phospho-null S325A mutant behaved as though no MITF were present. These data indicate that S325 phosphorylation, and the absence of K316 SUMOylation, delays HRR. We also transfected HA-MITF or the HA-MITF-S325A and S325E mutants into HEK293 cells and measured the accumulation of γH2AX after up to 6 h post-UV exposure (Supplemental Fig. S5A and 5B). As before, expression of HA-MITF triggered accumulation of γH2AX 2 h after UV irradiation. Expression of the phospho-mimetic S325E further enhanced the accumulation of γH2AX while the MITF S325A mutant was largely able to revert the effect of MITF. We performed a similar experiment using MITF E318K and determined that this mutant was able to activate γH2AX 4-times more than MITF WT (Extended Data Fig 5c and 5d).

**Figure 5.**
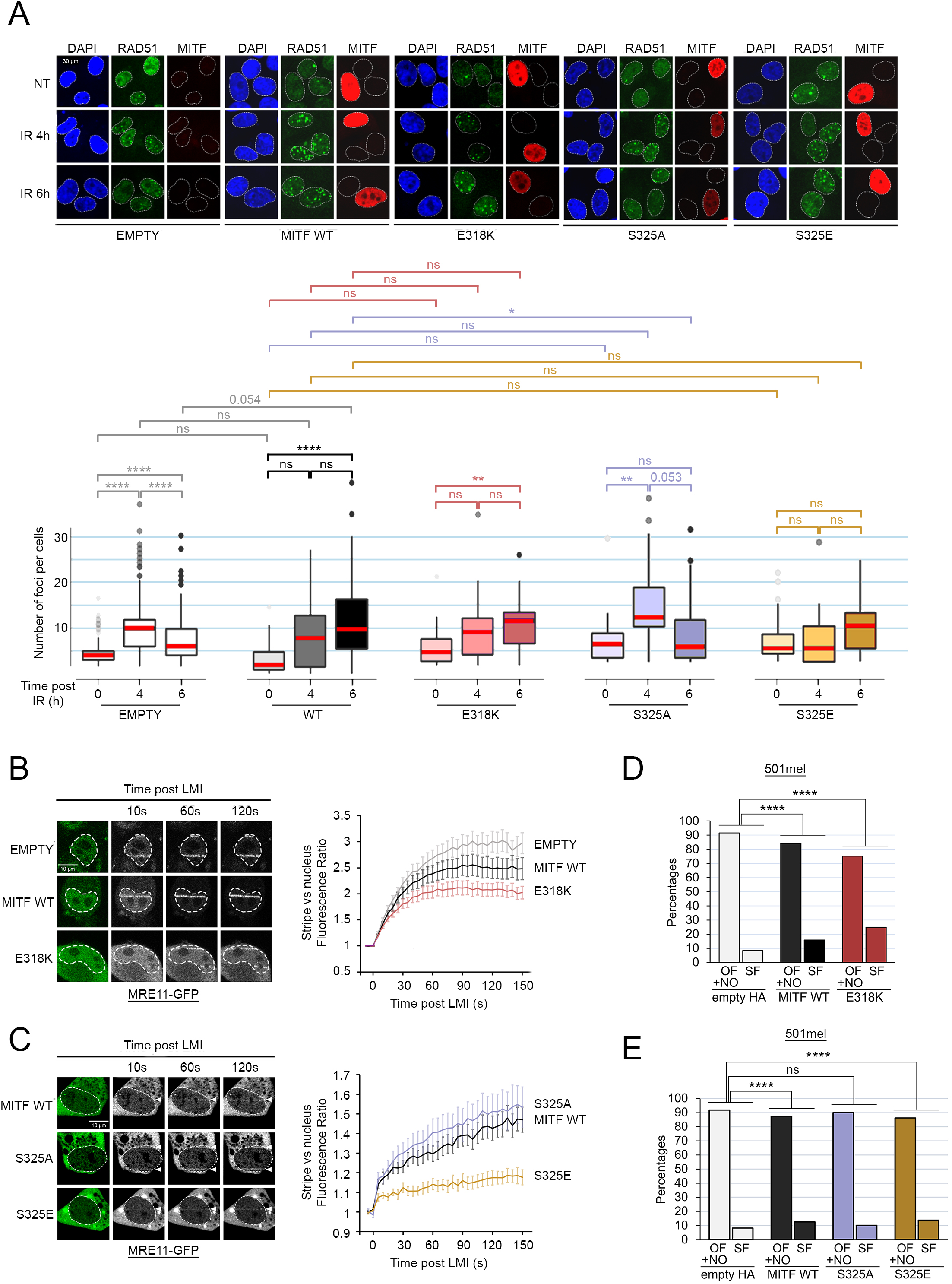
Effects of S325 and E318 mutations on MITF-mediated genome instability. (*A*) Immunofluorescence images (*top*) and quantification (*bottom*) of the formation of RAD51 foci after IR (X-rays, 2 Gy) in U2-OS cells transfected with HA-MITF WT (grey/black), S325A (purple), S235E (gold), or E318K (red) or the corresponding empty vector (white). Cells were fixed and processed at the indicated times after IR. The *p-values* were computed using the paired Wilcoxon Rank Sum test. *ns*: non-significant; *: *p<0.05;* **: *p<0.01;* ****: *p<0.0001*. The medians are indicated in red. (*B - left*) Still images of MRE11-GFP LMI in the presence of MITF WT, E318K mutants or the corresponding empty vector. Nuclei are delimited using the pre-irradiation images. The positions of the irradiated lines are indicated with an arrow. (*B - right*) Quantification MRE11-GFP recruitment. The graph represents the mean +/- SEM stripe/nucleus ratio over time. Values are normalized against the pre-LMI measurements. The baseline is indicated with a red dotted line. (*C - left*) Still images of MRE11-GFP LMI in the presence of MITF WT, S325A or S325E mutants. Nuclei are delimited using the pre-irradiation images. The positions of the irradiated lines are indicated with an arrow. (*C - right*) Quantification MRE11-GFP recruitment. The graph represents the mean +/- SEM stripe/nucleus ratio over time. Values are normalized against the pre-LMI measurements. The baseline is indicated with a red dotted line. (*D*) Graph expressing the percentage of ongoing forks (OF)/new origins (NO) and stalled replication forks (SF) in 501mel cells overexpressing MITF WT (black), E318K (red) or the corresponding empty vector (grey). *P-values* were determined using Fisher’s exact test. *p(MITF WT)*= 2.461506e-06; *p(E318K)*= 1.605343e-20. EMPTY HA and MITF WT are the same as in Fig. 1H. (*E*) Graph expressing the percentage of ongoing forks (OF)/new origins (NO) and stalled replication forks (SF) in 501mel cells overexpressing MITF WT (black), S325A (purple), S325E (gold) or the corresponding empty vector (grey). *P-values* were determined using Fisher’s exact test. *p(MITF WT)*= 6.942773e-07; *p(S325A)*= 6.332160e-02; *p(S325E)*= 2.172679e-07.

Having determined that MITF WT could block MRE11 recruitment to DNA damage, which would explain the deficiency in HRR and the delayed accumulation of RAD51 foci, we examined the effects of the MITF mutants. MRE11-GFP together with MITF WT, the E318K mutant, or the control HA-only vector, were expressed in U2-OS cells that were subject to LMI (Fig. 5B). We again observed a moderate reduction in MRE11 accumulation at DNA damage when MITF WT was expressed. Strikingly, this was more pronounced with the SUMOylation-defective mutant. We then repeated this assay using the S325A and S325E mutants. While in the presence of the phospho-null MITF-S325A MRE11 showed a slight increase in the accumulation to damage compared to MITF WT, the phospho-mimetic S325E largely blocked MRE11 recruitment (Fig. 5C). Using the nuclear tethering assay, we observed that MITF-NBS1 interaction is dependent on S325 status as seen with the low levels of colocalization between NBS1 and MITF-S325A (Supplemental Fig. S5E). At the same time, the S325E and E318K mutants retained the ability to bind NBS1, indicating that the MITF-NBS1 interaction may be involved in the inhibition of MRE11 recruitment. Taken together, our LMI data showed that the negative effect of MITF on MRE11 recruitment is dependent on S325 phosphorylation and potentiated by the SUMOylation-inhibiting E318K mutation.

Given these results, we would predict that the phosphorylation and SUMOylation mutants would affect HRR and MITF-induced replication stress. Using the U2-OS DR-GFP HRR reporter assay, cells were transfected with the I-SceI-expressing vector and co-transfected with either the empty HA vector, WT HA-MITF or the SUMOylation defective MITF-E318K mutant (Supplemental Fig. S5F). We confirmed the 20% reduction in HRR efficiency in the presence of HA-MITF compared to I-SceI alone and observed an equivalent reduction in HRR efficiency when expressing MITF-E318K. A BRCA2-specific siRNA was used as a control. We then performed the DNA fibre assays in 501mel cells expressing MITF WT, or either mutant. We confirmed that expression of MITF WT increased the percentage of stalled replication forks (Fig. 5D and 5E). Strikingly, MITF E318K expressing cells displayed an even higher proportion of stalled forks (Fig. 5D). On the other hand, cells expressing the S325E mutant displayed a proportion of stalled forks similar to that of MITF WT, while S325A-expressing cells showed no increase (Fig. 5E). Notably, MITF E318K-expressing cells have a slower replication speed overall which likely reflects the increase in stalled forks (Supplemental Fig. S4G). Overall, we conclude that the function of MITF in HRR and DNA replication are governed by post-translational modifications on K316 and S325.

In conclusion, we describe how in response to DNA damage, MITF interacts with the MRN complex and limits its recruitment to DNA, thus causing a defect in HR-mediated repair and an increase in replication stress in an ATM-dependent manner. We also demonstrate that the E318K germline mutation circumvents the need for S325 phosphorylation, while the MITF-E318K mutant can further increase the amount of replication stress. A model consistent with these results is presented in Fig. 6.

**Figure 6.**
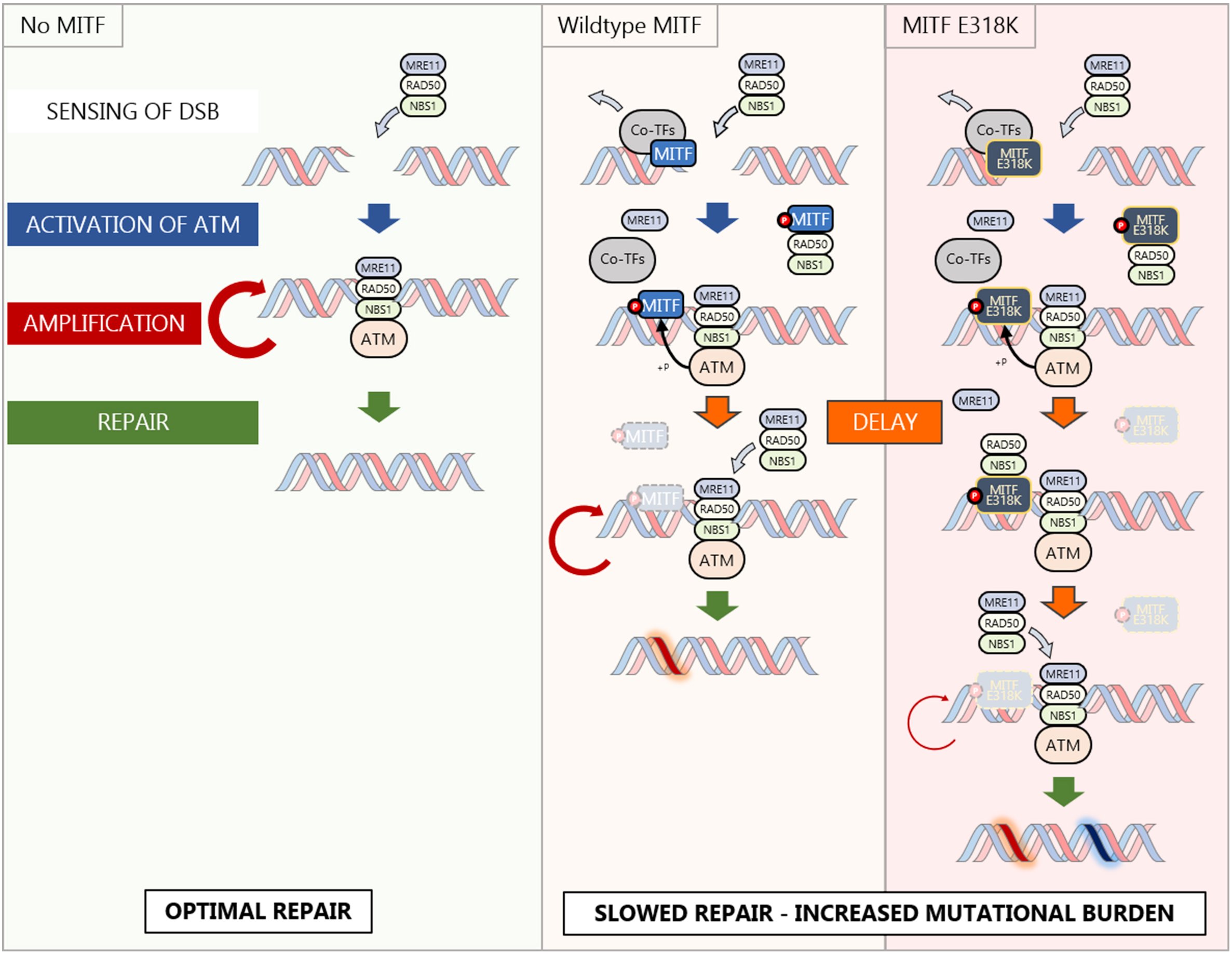
The impact of MITF expression on HR-mediated repair. In cells that do not express MITF (left panel), the MRN complex senses DNA damage, binds to DSBs, and then activates ATM to amplify the DDR and trigger repair by HRR. In melanocytes and melanoma cells (middle panel), the initial recruitment of MRN coincides with the detachment of MITF transcription co-factors. MITF is phosphorylated by ATM and DNA-PK and interacts with NBS1-RAD50 but not MRE11, destabilising the MRN complex. As a result, the rate of MRN recruitment is decreased, slowing down the DDR amplification step and the subsequent HR-mediated repair. Later, MITF is degraded, and the MRN complex is restored. The delay in MRN recruitment is emphasized in cells expressing the mutated MITF-E318K (right panel) because the resulting protein is more stable and more slowly degraded. The overall consequence of a delayed HRR is an increased risk of genome instability and mutational burden.

## Discussion

Maintenance of the integrity of the genome is regarded as one of the most important cellular functions, since failure of DNA repair mechanisms may lead to an increased mutation burden with major deleterious consequences including senescence or malignant transformation. On the other hand, under stress conditions, generation of genetic diversity arising from inefficient DNA damage repair may present an advantage by ensuring that some cells within a population may be better adapted to survive. Yet, while many of the mechanisms underpinning DNAdamage repair are common to all cells, some cell types exposed to distinct types of DNA damage or microenvironmental stresses, or located in specific anatomical positions, may superimpose lineage-restricted modulation of their DNA repair functions. In melanocytes and melanoma, MITF has received considerable attention as a critical coordinator of many cell functions (Goding and Arnheiter 2019). Previous work established a transcriptional role for MITF in regulating genes implicated in DDR (Strub et al. 2011), recapitulated here, as well as in promoting survival and pigmentation following UV exposure of the skin. MITF expression after UV irradiation exhibits a damped oscillation pattern (Malcov-Brog et al. 2018) in which it seems likely that a MITF negative feedback loop may participate (Louphrasitthiphol et al. 2019). However, although examination of SNVs in melanoma previously failed to identify any correlation between MITF expression and accumulated DNA damage (Herbert et al. 2019), here we revealed that both low and high levels of MITF correlated with an increased tumour mutational burden (TMB). While the detection of increased SNVs in MITF^Low^ cells might be explained by reduced expression of MITF target genes involved in DDR such as *BRCA1*, *LIG1*, *RPA1*, *FANCA* or *FANCC* (Strub et al. 2011), why MITF^High^ cells might also have a high TMB was not clear. We considered two possibilities. First, MITF^High^ cells may have a survival advantage that allows them to resist cell death following DNA damage leading to an increased TMB in the surviving cells. Consistent with this, MITF can activate expression of the anti-apoptotic effector *BCL2* within a few hours following UV irradiation (McGill et al. 2002; Malcov-Brog et al. 2018). A second, but not mutually exclusive, possibility is that cells expressing higher levels of MITF trigger replication stress that consequently leads to accumulation of DNA damage. The ability of MITF to cause replication stress may be related to its capacity to induce expression of pro-proliferative genes such as *CDK2* (Du et al. 2004), *MET* (McGill et al. 2006), *E2F1* (Chauhan et al. 2022) and *SCD* (Vivas-Garcia et al. 2020), and to suppress anti-proliferative factors such as p27 and p16 (Loercher et al. 2005; Carreira et al. 2006). The association between MITF expression and DNA replication in melanocytes is consistent with what is observed after UV irradiation in the skin. Following UV exposure, keratinocytes mediate the activation of MC1R (Guida et al. 2022) that is expressed in melanocytes. Signalling downstream from MC1R can promote DDR (Swope et al. 2014; Robles-Espinoza et al. 2016), but can also increase MITF levels (Khaled et al. 2003)that may subsequently lead first to increased proliferation and later increased pigmentation (Malcov-Brog et al. 2018).

Our results also indicate that following DNA damage, the MITF interactome is dramatically remodelled. MITF interaction with the majority of transcription co-factors was greatly reduced, with the notable exception of EP300 and CREBBP that can bind and acetylate MITF on multiple lysines (Louphrasitthiphol et al. 2020). We view it as likely that the loss of interaction with transcription cofactors arises following DNA damage as a consequence of the rapid degradation of the majority of MITF that is not retained at the DNA damage site; degradation MITF associated with transcription cofactors would explain loss of co-factor interaction. These observations suggest that the ability of MITF to regulate transcription may be impaired, at least at a subset of genes where specific co-factors may be required. At first sight, this may seem at odds with the known ability of MITF to activate transcription of survival, proliferation, and pigmentation gene programs after UV irradiation (Malcov-Brog et al. 2018). However, the transcriptional response of these MITF targets appears to occur 7-10 h after UV. It seems likely therefore that at early times (within minutes) DNA damage decreases MITF association with transcription co-factors, and the consequent reduction in MITF transcriptional output may suppress any MITF transcription-driven replication stress, but at later times (within hours) the ability of MITF to activate transcription of its downstream target genes may be restored.

While MITF loses its interaction with the majority of transcription co-factors following DNA damage, its association with the NBS1 and RAD50 components of the MRN complex is maintained. The most prevalent interaction was with NBS1, whose role is to facilitate the localisation of the MRN complex to DNA damage sites that contributes to the optimal activation of ATM (Uziel et al. 2003). We could detect the MITF/NBS1 interaction in steady state condition although it is stronger after camptothecin treatment. This suggests that MITF may titrate NBS1 away from the damage sites and from MRE11 and RAD50 and diminish the availability of the complex after DNA damage (see model in Fig. 6). Indeed, we observed that expression of MITF impacts the intensity of recruitment of the MRN complex at laser-induced double strand breaks. A couple of recent studies proposed an absolute quantitation of most proteins in human cells (Beck et al. 2011) or tissues (Jiang et al. 2020). In the report from Jiang *et al*. (Jiang et al. 2020), it was shown that in the skin NBS1 is the limiting factor amongst the MRN complex with an average of 784 copies of NBS1, 1345 copies of MRE11 and 5000 copies of RAD50. Beck *et al*. (Beck et al. 2011) measured the proteome of U2-OS cells and quantified between 5480 and 8030 copies of MRN proteins. By contrast, in 501mel cells, a high MITF-expressing cell line, the number of MITF dimers is estimated at up to 250,000 copies (Louphrasitthiphol et al. 2020). Thus, titration of NBS1 by an excess of MITF is feasible and would have an effect on the recruitment of all three proteins. As a consequence, MITF may limit the recruitment of the MRN complex to sites of damage leading to reduced HRR, increased replication stress and higher TMB. By limiting MRN activity, MITF may create a tolerance to DNA damage that may be important to allow cells to survive, especially in the case of prolonged sun exposure when MITF-dependent melanocyte proliferation and melanin production are essential (Goding 2007). The ability of MITF to influence the DDR resembles ATF2 which is also found at damage sites. However, in contrast to MITF, ATF2 enhances the recruitment of the MRN complex (Bhoumik et al. 2005). Interestingly, ATF2 is a co-activator of Jun whose transcriptional program is associated with melanoma dedifferentiation and low MITF (Riesenberg et al. 2015; Verfaillie et al. 2015; Comandante-Lou et al. 2022), raising the possibility that the DDR is affected differently in distinct phenotypic states.

Unexpectedly, MITF is retained at damage sites as consequence of a retention mechanism controlled by ATM and DNA-PK-dependent phosphorylation and is inhibited by SUMOylation, while in the rest of the nucleus MITF levels decrease. This is fundamentally different from active recruitment as observed with *bona fide* repair proteins like NBS1 or MRE11 and may indicate that MITF is a modulator of the DDR rather than an essential player. Interestingly, SUMOylation and by extension the E318K mutation seem to affect both the behaviour of MITF at, and away from, the damage sites. Indeed, only the MITF-E318K mutant showed an increase at the laser-induced DNA damage sites, suggesting its active recruitment in addition to retention. Moreover, the overall decrease in MITF-E318K was much slower than the WT protein or the phosphorylation mutants. Because MITF-E318K is also able to interact with NBS1, disrupting the MRN complex and preventing MRE11 recruitment and function, we observe an even higher accumulation of γH2AX due to a delayed HRR in MITF-E318K expressing cells that also have a higher percentage of stalled replication forks and slower fork speed, and resist senescence-induced terminal growth arrest (Bonet et al. 2017; Leclerc et al. 2017). This combination of delayed HRR and tolerance to replication stress may therefore explain the increased melanoma risk associated with the MITF-E318K mutation. Importantly, we also revealed that the S325E phospho-mimetic phenocopies the E318K mutation, that inhibits SUMOylation at K316, with both accumulating more at sites of DNA damage and both able to bind NBS1 and reduce replication fork speed.

MITF is a tissue-restricted transcription factor, and we reveal that it can play a non-transcriptional role in the DDR. It is likely that MITF is not unique in this respect and that more transcription factors regulate the DDR in a context-specific manner. Here, the MITF-dependent regulation of HRR is mediated by the interaction with NBS1 and regulated by ATM and DNA-PK kinases. Although several transcription factors can be phosphorylated by DDR-associated kinases or interact with DDR factors, only a few have been implicated in DNA repair (Bhoumik et al. 2005; Rubin et al. 2007; Beishline et al. 2012; Park et al. 2015; Wysokinski et al. 2015), and a study from Izhar *et al*. (Izhar et al. 2015) suggested that the recruitment of transcription factors to DNA damage was an artefact of their affinity for PARP1. Here instead, we showed that MITF retention at sites of damage is not only regulated, but that its effects on replication fork progression and DNA damage repair are mediated though a specific interaction with NBS1. Thus, we propose that the transcription factor-dependent regulation of the DDR may be a general mechanism that allows fine tuning of the repair processes depending on the tissue or the context.

In summary, our results provide a key insight into how a tissue-restricted transcription regulator and lineage survival oncogene, MITF, can shape the response to DNA damage, and may provide a potential mechanistic explanation for how the E318K mutation impacts melanoma predisposition.

## Materials and Methods

### SNV count

Gene expression, simple somatic mutation, and donor data have been downloaded from the International Cancer Genome Consortium (ICGC) data portal for the Skin Cutaneous melanoma - ICGC project (SKCM-US) from https://dcc.icgc.org/releases/current/Projects/SKCM-US. The 428 samples were sorted by their MITF expression value and split into 5 bins of approximately equal size. The distribution of single nucleotide variants (SNVs) per sample were plotted after log-transformation computed as log10(0.1+SNV). Paired Wilcoxon Rank Sum test was used to compare the values between the bins.

### Melanoblast/melanocytes differentiation protocol

The exact procedure to differentiate melanoblasts and melanocytes from hPSCs was described previously (Baggiolini et al. 2021).

### DNA extraction

DNA from frozen cells was isolated with the DNeasy Blood & Tissue Kit (QIAGEN catalog # 69504) according to the manufacturer’s protocol modified by replacing AW2 buffer with 80% ethanol. DNA was eluted in 50 µL 0.5X Buffer AE heated to 55°C.

### IMPACT (Cheng et al. 2015)

After PicoGreen quantification and quality control by Agilent BioAnalyzer, 100 ng of DNA were used to prepare libraries using the KAPA Hyper Prep Kit (Kapa Biosystems KK8504) with 8 cycles of PCR. 200-400 ng of each barcoded library were captured by hybridization in a pool of 5 samples using the IMPACT (Integrated Mutation Profiling of Actionable Cancer Targets) assay (Nimblegen SeqCap), designed to capture all protein-coding exons and select introns of 468 commonly implicated oncogenes, tumor suppressor genes, and members of pathways deemed actionable by targeted therapies. Captured pools were sequenced on a HiSeq 4000 in a PE100 run using the HiSeq 3000/4000 SBS Kit (Illumina) producing an average of 866X coverage per tumor and 538X per normal. Sequence reads were aligned to human genome (hg19) using BWA MEM (Li and Durbin 2010). ABRA was used to realign reads around indels to reduce alignment artifacts, and the Genome Analysis Toolkit was used to recalibrate base quality scores (McKenna et al. 2010; Mose et al. 2014). Duplicate reads were marked for removal, and the resulting BAM files were used for subsequent analysis. The sequencing data analysis pipeline can be found here: https://github.com/soccin/BIC-variants_pipeline. Copy number analysis was performed using FACETS (Shen and Seshan 2016) (version 0.3.9) using a set of pooled normals for each cell line.

### Cell culture and drugs

All cell lines were grown in DMEM containing 10% FBS at 37ᵒC and 5% CO_2_. U2-OS-DR-GFP, 501-HA-MITF wildtype and mutants and U2OS-LacO#13 were maintained under puromycin selection. MITF-GFP (wild type and mutants) expressing 501mel were selected and maintained in geneticin. HEK293 cells were transfected with Fugene 6 (Promega). All other cell lines were transfected using Lipofectamine 2000 (ThermoFisher Scientific). All transfections were performed following the manufacturer’s instructions. Doxycycline was used for all inducible cell lines.

| Drug/Chemical | Supplier | Reference |
| --- | --- | --- |
| Doxycycline | Sigma-Aldrich | D9891 |
| Puromycin | Sigma-Aldrich | P8833 |
| Geneticin | ThermoFischer Scientific | 10131027 |
| KU55933 | Tocris | 3544 |
| VE821 | Selleckchem | S8007 |
| VE822 | Selleckchem | S7102 |
| NU7441 | Selleckchem | S2638 |
| Olaparib | Selleckchem | S1060 |
| Camptothecin | Cambridge Bioscience | 1039-1 |
| CHK2 purified kinase | Life Technologies | PV3367 |
| DNAPKcs purified kinase | Life Technologies | PV5866 |
| Cycloheximide | Merck | C4859 |

### Plasmids and oligonucleotides

All plasmids were described elsewhere (Table). Point mutations were performed using the QuikChange Lightning site-directed mutagenesis kit (Agilent Technologies) according to manufacturer’s instructions. Mutagenesis primers (Table) were designed using the Agilent online tool (https://www.agilent.com/store/primerDesignProgram.jsp). For MITF silencing, we used a previously validated sequence (Carreira et al. 2006) and transfected cells using Lipofectamine RNAiMax (ThermoFisher Scientific) as per the manufacturer’s instructions.

| Plasmid | Source |
| --- | --- |
| pCMV-3XHA | Halazonetis |
| pEGFP-C1-MITF | (Ngeow et al. 2018) Ngeow *et al.*, 2018 |
| RFP-SUMO1/2 WT and mutants | Jo Morris, Birmingham |
| GFP-MRE11 | Tim Humphrey |
| mCherry-LacR-NLS | Greenberg |
| I-SceI | Tim Humphrey |
| pcDNA3.1(+)-C-eGFP-NBS1 | GenScript |
| pcDNA3.1(+)-C-eGFP-RAD50 | GenScript |
| pLUC-Met | Generated in our lab |

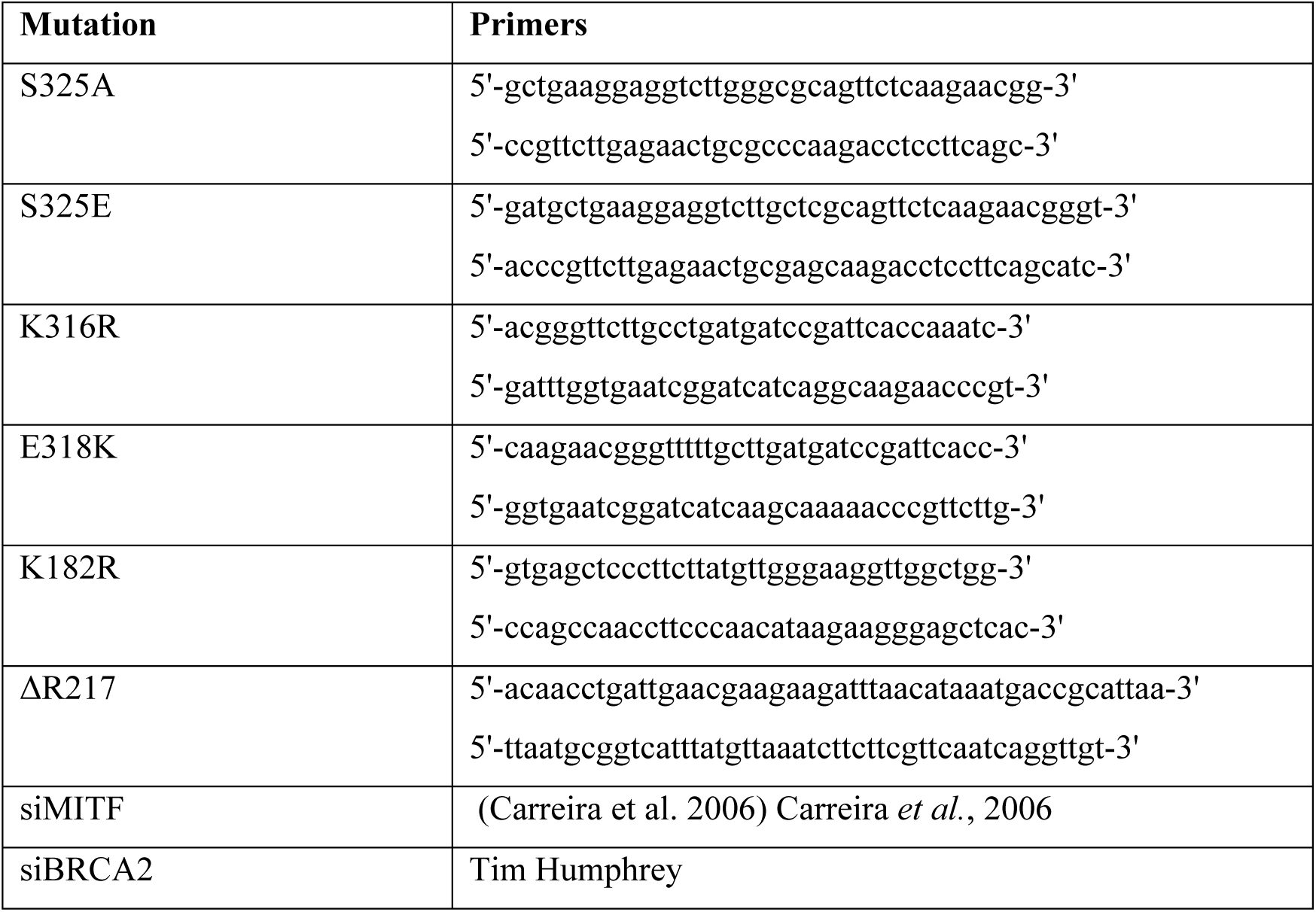

### Peptide arrays for kinase assays

Cellulose-bound peptide arrays were prepared using standard Fmoc solid phase peptide synthesis on a MultiPep-RSi-Spotter (INTAVIS, Köln, Germany) according to the SPOT synthesis method provided by the manufacturer, as previously described (Picaud and Filippakopoulos 2015). In brief, twenty-one-residue-long peptides based on MITF residues 314 to 333 were synthesized on amino-functionalized cellulose membranes (Whatman Chromatography paper Grade 1CHR, GE Healthcare Life Sciences #3001-878) and the presence of SPOTed peptides was confirmed by ultraviolet light (UV, λ = 280 nM). The wild-type sequence was mutated or modified where indicated. The membrane was quickly washed in EtOH 100% then equilibrated overnight in Kinase Buffer (HEPES 50 mM pH7.5, NaCl 200 mM, MgCl2 10 mM, KCl 10mM). For *in vitro* kinase assays, the membrane was then incubated in Kinase Buffer containing 500 ng of kinase for 1 h at 30 °C. After washing, the membranes were analysed by western blot. The wildtype sequence was mutated or modified where indicated. The membrane was briefly incubated in EtOH 100% followed by an overnight incubation in the Kinase Buffer (HEPES 50mM ph7.5, NaCl 200 mM, MgCl_2_ 10 mM, KCl 10mM). For the *in vitro* kinase assays, the membrane is then incubated in the Kinase Buffer containing 500 ng of the kinase for 1 h at 30°C. After washing, the membrane is analysed by western blot.

### Western blot and antibodies

Cells were lysed with 1X Laemmli sample buffer (50 mM Tris-Cl (pH6.8), 100 mM DTT, 2% SDS, 12.5% glycerol, 0.1% Bromophenol blue), sonicated in a Diagenode Bioruptor (4ᵒC, 30 s ON/15 s OFF cycles, 5 min) and boiled at 95ᵒC for 15 min. Protein separation was performed by SDS–polyacrylamide gel electrophoresis (SDS-PAGE) using 12.5% 200:1 bis-acrylamide gels. Proteins were transferred onto nitrocellulose membranes and saturated in PBS-BSA 3% for 1 h. The membranes were incubated with the primary antibody at 1ug/mL in PBS-BSA 1% overnight, followed by 1 h incubation with the HRP-conjugated secondary antibody diluted 1/10,000 (mouse) or 1/20,000 (rabbit) times in PBS-Tween 0.05%. Detection of the chemo-luminescent signal was performed using the Amersham ECL Western Blot Detection (GE Healthcare) according to the manufacturer’s instructions. Images were acquired on a Bio-Rad ChemiDoc™ XRS+ System and analysed with the Image Lab software.

| Target | Supplier | Reference |
| --- | --- | --- |
| 53BP1 | Cell Signaling Technology | 4937 |
| FLAG | Merck (Sigma-Aldrich) | F1804 |
| GAPDH | Merck (Sigma-Aldrich) | G9545 |
| GFP | Abcam | ab290 |
| HA Tag | Merck (Sigma-Aldrich) | H3663 |
| HA Tag | Cell Signaling Technology | 3724 |
| MITF | Cambridge Bioscience | HPA003259 |
| MITF D5 | Abcam | ab3201 |
| MITF phospho-S325 | Custom-made |  |
| Phospho-ATM (Ser1981) | Abcam | ab81292 |
| Phospho-H2AX (Ser139) | Merck (Millipore) | 05-636 |
| Phospho-H2AX (Ser139) | Cell Signaling Technology | 9718 |
| RAD51 | Cell Signaling Technology | 8875 |
| RFP | Abcam | ab152123 |
| Anti-Mouse HRP | Bio-Rad | 1706516 |
| Anti-Rabbit HRP | Bio-Rad | 1706515 |
| Anti-Mouse Alexa Fluor 488 | ThermoFisher Scientific | A11001 |
| Anti-Mouse Alexa Fluor 546 | ThermoFisher Scientific | A10036 |
| Anti-Rabbit Alexa Fluor 488 | ThermoFisher Scientific | A11008 |
| Anti-Rabbit Alexa Fluor 546 | ThermoFisher Scientific | A10040 |

### Immunofluorescence

For immunofluorescence-based experiments, cells were seeded on coverslips or in 8-chamber glass-bottom slides (Ibidi). Cells were fixed in 4% formaldehyde and permeabilised in 0.1% Triton X-100/1xPBS. For LMI or foci experiments, cells were first incubated in the CSK-T buffer (PIPES 10mM pH7, NaCl 100 mM, Sucrose 300 mM, MgCl_2_ 3 mM, 0.5% Triton X-100) then fixed in formaldehyde. Samples were saturated in PBS-BSA 3% for 1 h, then incubated with the primary antibody at in PBS-BSA 1% from 1h to overnight, followed by 1h incubation with the fluorochrome-conjugated secondary antibody. When required, the nuclear content was counterstained with DAPI or Hoechst. Coverslips were mounted on glass slides using VECTASHIELD® Antifade Mounting Medium (Vectorlabs). When 8-chambers slides were used, samples were covered with 9 mm coverslips. Analysis of the fluorescent patterns was performed on a Zeiss LSM710 confocal microscope. Quantification of the DDR foci was automatized using the ImageJ2 software and self-created macros.

### Laser Microirradiation

For live cells LMI, cells were cultured in µ-Dish 35 mm (Ibidi #81158). 24h after transfections, cells were pre-sensitised with 1ug/mL of Hoechst 33342 for 30 min. Medium was immediately replaced with FluoroBrite DMEM (ThermoFisher Scientific) supplemented with FBS and glutamine. For UV-LMI, UV irradiation was performed on a Nikon TE-2000 with Nikon Plan Fluor 20x/0.45 (Nikon, Surrey, UK) using a Team Photonic SNV-04P-100 laser (Cairn Research, Kent, UK), a plan-apochromat 20X/0.8 M27 objective and a Prime camera (Photometrics, Arizona, US). For NIR-LMI, Irradiation was performed on a Zeiss LSM710 confocal microscope using a MaiTai multiphoton laser (Spectra Physics) and a plan-apochromat 63X/1.40 oil DICM27 objective. We used a 750nm laser which at 100% provides 0.5W at the objective. Cells were exposed to 5% to 12% power depending on the application. Analysis and quantification of the fluorescent stripes was automatized using the ImageJ2 software and self-created macros. When the samples were destined to IF, cells were plated on large coverslips, pre-sensitised and irradiated through a plan-apochromat 20X/0.8 M27 objective, further incubated for the required time and processed as indicated.

### Nuclear tethering

U2OS-LacO#13 cells (Lu et al. 2021) were grown on coverslips and transfected with the indicated combinations of plasmids for 24 h. Samples were fixed with formaldehyde and mounted as previously indicated. Single cells images were acquired using a Zeiss LSM710 confocal microscope and the intensities of colocalization was determined manually.

### DNA Fibre Assay

Cells were incubated with 10 μM BrdU for 30min followed by 10 μM EdU for 30min and collected. The DNA fibre spreads were prepared as recommended (Nieminuszczy et al. 2016). After DNA denaturation and blocking step, EdU was detected using the Click-iT® EdU Imaging Kit 488 (ThermoFischer Scientific) using 250 μL of Click-It reaction mix per slide for 2h. After another blocking step, BrdU was detected with a mouse anti-BrdU antibody clone 3D4 (BD Bioscience, #555627) and the anti-mouse Alexa Fluor 546. Images were acquired using a Zeiss LSM710 confocal microscope. Double-labelled replication forks were analysed manually using ImageJ2. For each sample, 300 to 400 were recorded. To determine the forks speed, the length of the EdU part of the ongoing forks was measured and we used the formula *V(kb/min) = [(x * 0.132 µm) * 2.59 kb/µm] / t (min)*, where *x* = length of EdU.

### Gamma and UV irradiation

For gamma irradiation, cells were exposed to X-rays at a final dose of 1 to 10 Gy using a cesium-137 irradiator at the dose rate of 1.87 Gy/min. For UV, the medium was replaced for PBS and plates were placed in a CL-1000 UV crosslinker containing 254nM tubes. After exposure to a final dose of 24 to 100J/m^2^, the culture medium was added back, and the cells were cultivated for the time required.

### DSB repair reporter assays

The procedure for the HR, SSA or BIR-DR-GFP reporter assay was described elsewhere (Ahrabi et al. 2016). U2-OS-DR-GFP cells were co-transfected with an I-SceI plasmid plus the indicated expression vector or siRNA using Lipofectamine 2000 (Invitrogen) for 72 h. The cells were trypsinised, washed and resuspended in cold PBS before flow cytometry analysis using a BD FACScanto™ (BD Bioscience). Results were generated using the FlowJo™ software (BD Bioscience).

### Proximity-Dependent Biotinylation Mass Spectrometry

The methods for the generation of MITF tagged cell lines and the proximity-dependent biotinylation mass spectrometry was described in detail elsewhere (Chauhan et al. 2022). Briefly, HEK293 cells stably expressing BirA*-FLAG-MITF or BirA*-FLAG-NLS were pelleted from two 150-mm plates. To induce DNA damage, cells were treated with 20 nM CPT for 4 h. After extraction, biotinylated proteins were captured using streptavidin-Sepharose beads for 3 h at 4⁰C, washed and recovered by incubating in a trypsin solution overnight at 37⁰C. The tryptic peptides were then stored at −80⁰C. For Mass Spectrometry analysis, eluted samples were analyzed on a TripleTOF 5600 instrument (AB SCIEX, Concord, Ontario, Canada) set to data dependent acquisition (DDA) mode. MS data storage, handling and analysis is detailed in the study by Chauhan et al 2022. For statistical scoring, we used Significance Analysis of INTeractome (Teo et al. 2014); SAINTexpress version 3.6.1) to model the background proteins observed in our BioID analysis and enforced a 1% false discovery rate threshold to identify significant proximity partners of MITF.

The MS files used in this report were deposited to MassIVE (http://massive.ucsd.edu) - accession number MSV000089109, and ProteomeXchange (http://www.proteomexchange.org/), accession number PXD032772. They can be found at ftp://MSV000089109@massive.ucsd.edu, using the usernames “MSV000089109_reviewer” and password “MITF”.

### Data

RNAseq data have been previously published (Louphrasitthiphol et al. 2019). MS data are available at MassIVE (http://massive.ucsd.edu; MSV000089109; ftp://massive.ucsd.edu/MSV000089109) and the ProteomeX-change Consortium (http://proteomecentral.proteomexchange.org; PXD032772).

## Acknowledgements

The authors would like to acknowledge Dr Jo Morris and Dr Alexander Garvin (Institute of Cancer and Genomics Sciences, University of Birmingham) for the SUMO vectors.

This work was funded by the Ludwig Institute for Cancer Research (CRG, RB, PL, MT), GABBA (DD), Marie Curie Cancer Care (SC), JPL was supported by a Junior 2 salary award from the Fonds de Recherche du Québec-Santé and an operating grant from the Cancer Research Society and Genome Quebec (25123). PF and SP were supported by the MRC: Medical Research Council (MR/N010051/1). SS and TH are funded by an MRC Programme Grant [MR/X006778/1 to T.C.H]. AB was supported by the Swiss National Science Foundation Postdoc mobility fellowship P2ZHP3_171967 and P400PB_180672. RW was supported by the NIH Director’s New Innovator Award DP2CA186572.

## Author Contributions

RB and CRG conceived the experiments. JPL performed the BioID analysis. SP and PF prepared the peptide arrays. DD performed the FACS experiments. MT performed the SNV analysis. RW, AB and SH performed the hPSC-based experiments. PL provided the RNAseq data and analysis. SS performed the gamma-irradiation based experiments. SC performed preliminary experiments. TH and CRG provided supervision and funding, and RB and CRG wrote the manuscript. All authors discussed the experiments and reviewed the manuscript.

## Author information

Correspondence and requests for material should be sent to CRG.

