## Supplemental methods and figures for "DNA damage-induced interaction between a lineage addiction oncogenic transcription factor and the MRN complex shapes a tissue-specific DNA Damage Response and cancer predisposition"

### SUPPLEMENTAL MATERIALS TABLE OF CONTENTS

**Supplemental Materials and Methods**

**Supplemental Figures and Movie List**

**Supplemental Figure Legends**

**Supplemental References**

### SUPPLEMENTAL MATERIALS AND METHODS

*Colony formation assay*

Cells were transfected in 6-wells plates. After 24 h, cells were counted and plated at low density (500, 1,000 or 2,000 cells per well) in a separate 6-well plate and let to grow for 10 days. Newly formed colonies were fixed in PFA, stained with Crystal Violet and dried. Plates were scanned using a FLA-5100 Fluorescent Image Analyzer (Fujifilm) and colonies manually counted.

*Luciferase reporter*

HEK293 cells were seeded in 12-wells plates and co-transfected with the MET-Luc plasmid and HA-MITF WT or mutant or the empty-HA vector. After 48 h, measurement of Luciferase activity was performed using the Dual-Glo® Luciferase Assay System (Promega) according to the manufacturer's instructions in a GloMax® Multi Detection System (Promega).

*RNAseq*

RNA was extracted using the RNeasy kit (Qiagen, 74106) and QC on a Bioanalyzer. Samples with RIN ≥9.5 were used for library prep and sequencing using Wellcome Trust genomic service, Oxford as previously described (Louphrasitthiphol et al. 2019). Briefly, siMITF library were prepared using the QuantSeq Forward kit (Lexogen, 015.96) using 500 ng of starting material to minimize the PCR amplification step with ERCC ExFold RNA spike-in mixes (Ambion) and sequenced on HiSeq 4000. The output raw fastq files were trimmed of poly-A using cutadapt (Martin 2011) and mapped using STAR 2.5.1b (Dobin et al. 2013) against hg38 (GRCh38, 2015). Counts per gene from STARwere used as input for differential gene expression analysis using EdgeR (Robinson et al. 2010). Reads for each sample set were first filtered for genes whose expression is less than one count per million. GSVA analyses were performed using the Bioconductor package GSVA (Hanzelmann et al. 2013). The GSVA matrices and gene specific expression table were clustered and displayed as a heat map using Pheatmap (https://cran.r-project.org/web/packages/pheatmap/index.html).

**SUPPLEMENTAL FIGURES**

Figure S1

Figure S2

Figure S3

Figure S4

Figure S5

**SUPPLEMENTAL MOVIES**

(Movie S1) GFP-MRE11 recruitment after LMI, in the presence or absence of MITF.

(Movie S2) GFP-NBS1 recruitment after LMI, in the presence or absence of MITF.

(Movie S3) GFP-RAD50 recruitment after LMI, in the presence or absence of MITF.

(Movie S4) GFP-MITF WT recruitment after LMI and treatment with DNA-PK inhibitor.

(Movie S5) GFP-MITF WT recruitment after LMI and treatment with ATM and ATR inhibitors.

(Movie S6) GFP-MITF WT recruitment after LMI and treatment with olaparib.

(Movie S7) GFP-MITF S325 and E318 mutants recruitment after LMI.

(Movie S8) GFP-MITF S325E recruitment after LMI and treatment with inhibitors.

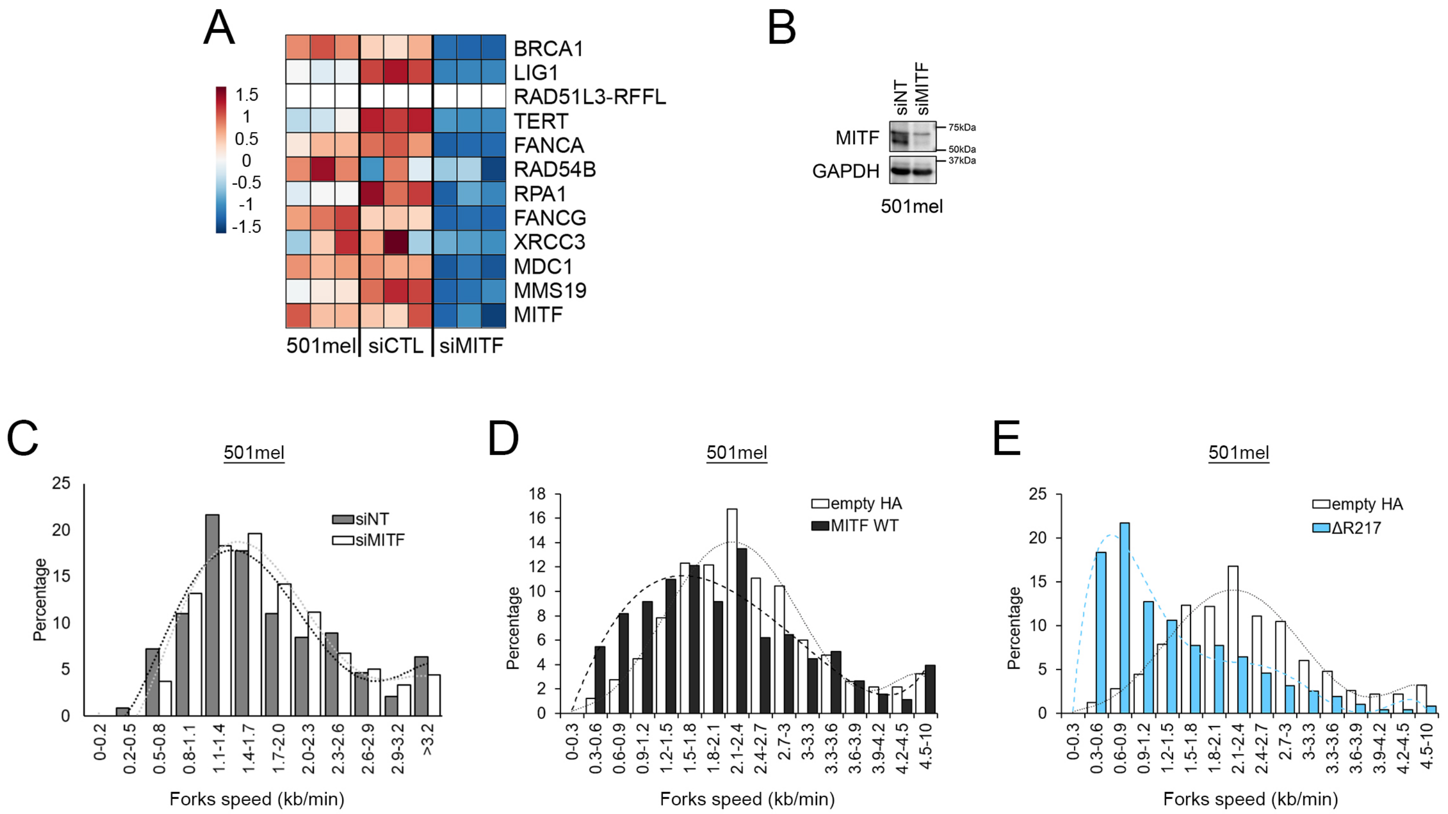


Binet et al. Supplemental Figure 1

**Figure S1.** (*A*) Heatmap derived from triplicate 3′ RNA-seq of 501mel cells control or transfected with siMITF or a non-targeting siRNA (siCTL) as indicated. The genes list is derived from Strub *et al.* (2011) (Strub et al. 2011). (*B*) Western blot of 501mel cells transfected with siMITF or a non-targeting control. GAPDH was used as loading control. (*C to E*). Histograms representing the distribution of the calculated fork speed in kilobase per minute in 501mel cells transfected with (*C*) siMITF (dark grey) or the non-targeting siRNA (white) and in 501mel cells transfected with (*D*) HA-MITF (black), (*E*) HA-MITF-delR217 (blue) or the corresponding empty vector (white). The dashed lines show the trendline for the matching bar graph.

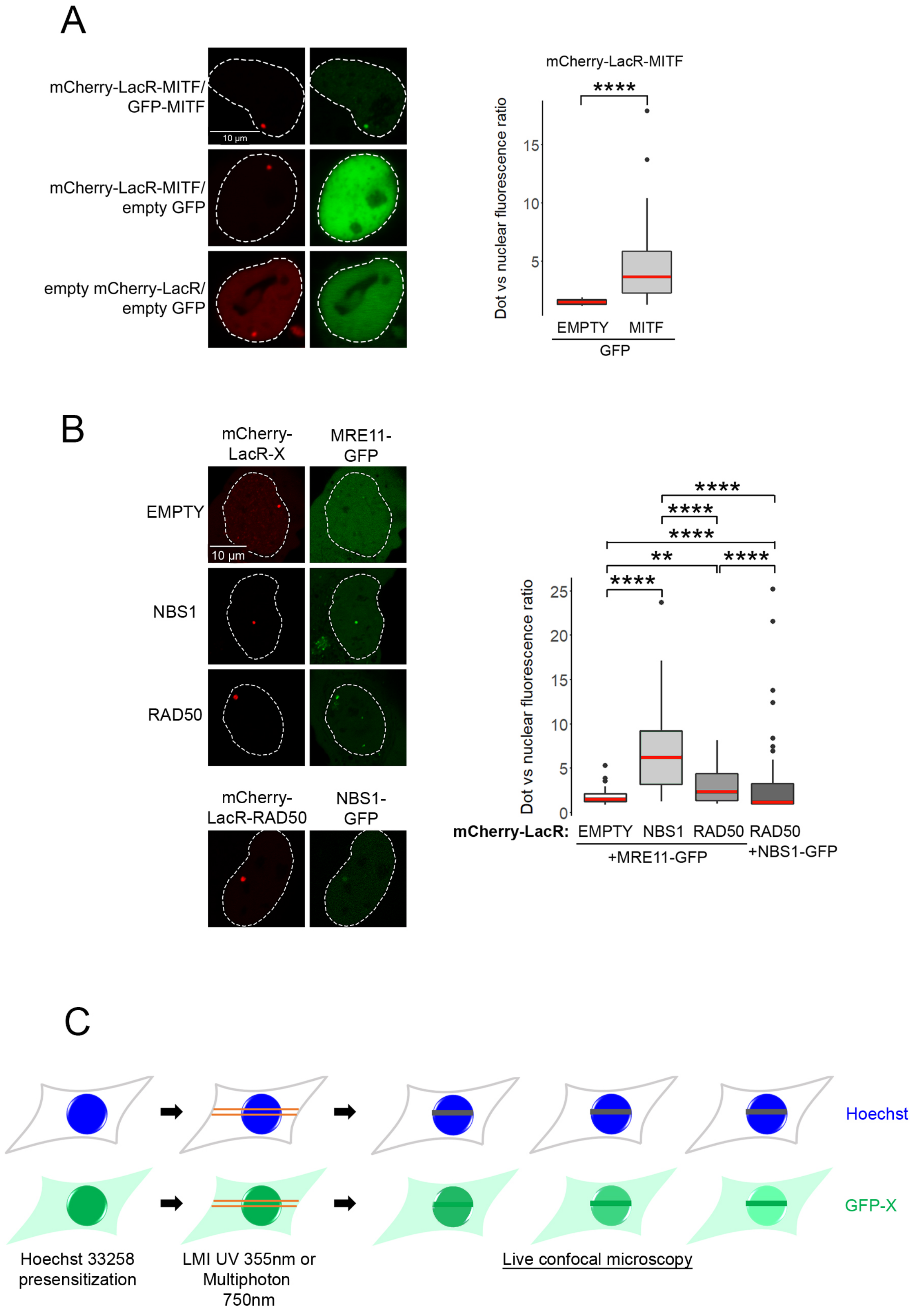


Binet et al. Supplemental Figure 2

**Figure S2.** (*A*) Representative images and quantification of the nuclear tethering assay showing dimerization of MITF. The left panels show the localization of mCherry-LacR-NLS or mCherry-LacR-MITF dots in the nuclei of U2OS-LacO#13 cells, and the right panels show GFP-MITF or GFP only. The quantification is expressed as the ratio between the fluorescence measured inside the mCherry dot and in the rest of the nucleus. The *p-values* were computed using the Wilcoxon Rank Sum test. ****: $p<0.0001$. The medians are indicated in red. (*B*) Representative images and quantification of the nuclear tethering assay showing interaction between MRE11 and RAD50 or MRE11 and NBS1. The left panels show the localization of mCherry-LacR-NLS, mCherry-LacR-NBS1 or mCherry-LacR-RAD50 dots in the nuclei of U2OS-LacO#13 cells, and the right panels show GFP-MRE11. The quantification is expressed as the ratio between the GFP fluorescence measured inside the area delimited by the mCherry dot and in the rest of the nucleus. The *p-values* were computed using the Wilcoxon Rank Sum test. **: $p<0.01$; ****: $p<0.0001$. The medians are indicated in red. (*C*) Principle of the LMI technique. Cells are presensitised with a DNA intercalant (Hoechst). Using a UV laser or a multiphoton laser, a limited area of the cells is irradiated resulting in the bleaching of the Hoechst dye. The recruitment of the protein of interest can be analysed by immunofluorescence or live video-microscopy.

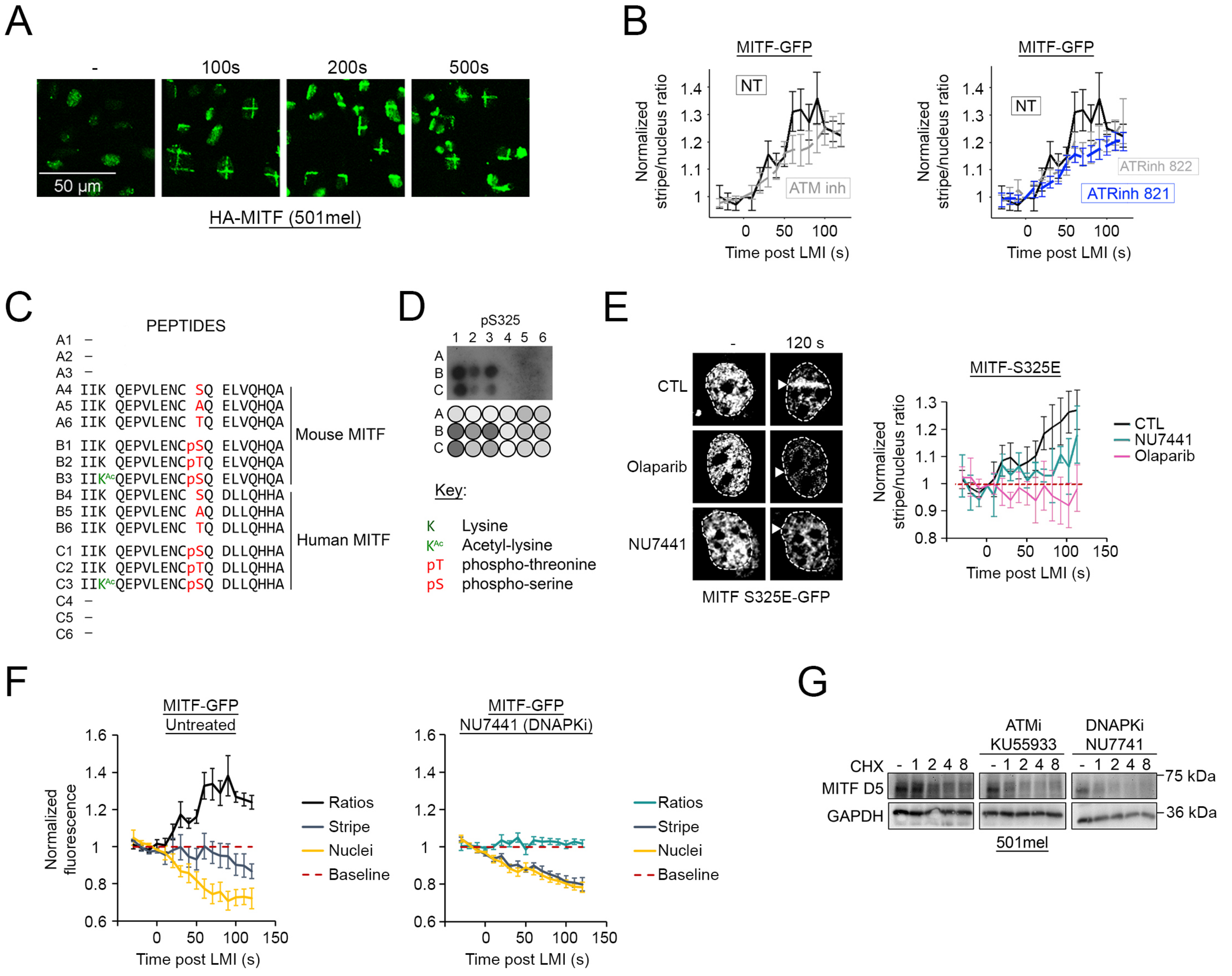


Binet et al. Supplemental Figure 3

**Figure S3.** (*A*) Immunofluorescence of stable 501HA-MITF cells after UV-LMI. The irradiated cells were fixed before UV-LMI or 100 s, 200 s and 500 s after UV-LMI and MITF detected with an anti-HA antibody. (*B*) Quantification of live video microscopy showing recruitment of MITF in 501mel cells stably expressing GFP-MITF after NIR-LMI. Cells were treated with ATM (KU55933 – 10 µM) or ATR (VE821 or VE822 – 1 µM) inhibitors for 24h before irradiation. The graphs represent the mean +/- SEM stripe/nucleus ratio over time. Values are normalized against the pre-LMI measurements. For clarity, each individual graph represents the same control curve (black) against only one type of inhibitor (in grey or blue). (*C and D*)**.** Validation of anti-phospho-S325 MITF antibody using peptide arrays. (*C*) Sequences of the peptides adsorbed in the array. (*D*) *Top:* Map of the peptide array and result of the western blot. *Bottom:* Key to the different symbols used in the list in A. (*E*) Still images and quantification of live video microscopy showing recruitment of MITF.S325E in 501mel cells stably expressing GFP-MITF after NIR-LMI. Cells were treated with DNA-PK (NU7441 - 1 µM) or PARP (Olaparib – 10 µM) inhibitors for 24h before irradiation. The graphs represent the mean +/- SEM stripe/nucleus ratio over time. Values are normalized against the pre-LMI measurements. (DNAPKi is in turquoise and PARPi is in pink). The baseline is indicated with a red dotted line. (*F*) Details of GFP-MITF behaviour in stable 501GFP-MITF cell lines after LMI, as known in Fig. 4c. Quantification of GFP intensities at the LMI sites (stripes – blue curves) and away from the LMI sites (nuclei – yellow curves). The graph represents the mean +/- SEM GFP fluorescence over time. Values are normalized against the pre-LMI measurements. The baselines are indicated with red dotted lines. (*G*) Time course of endogenous MITF expression levels after CHX treatment. Western blot of 501mel cells treated with CHX and harvested at the indicated time points. Cells were either mock-treated or exposed to the indicated inhibitors (KU55933 – 10 µM, NU7441 – 1 µM) for 16 h. GAPDH was used as loading control.

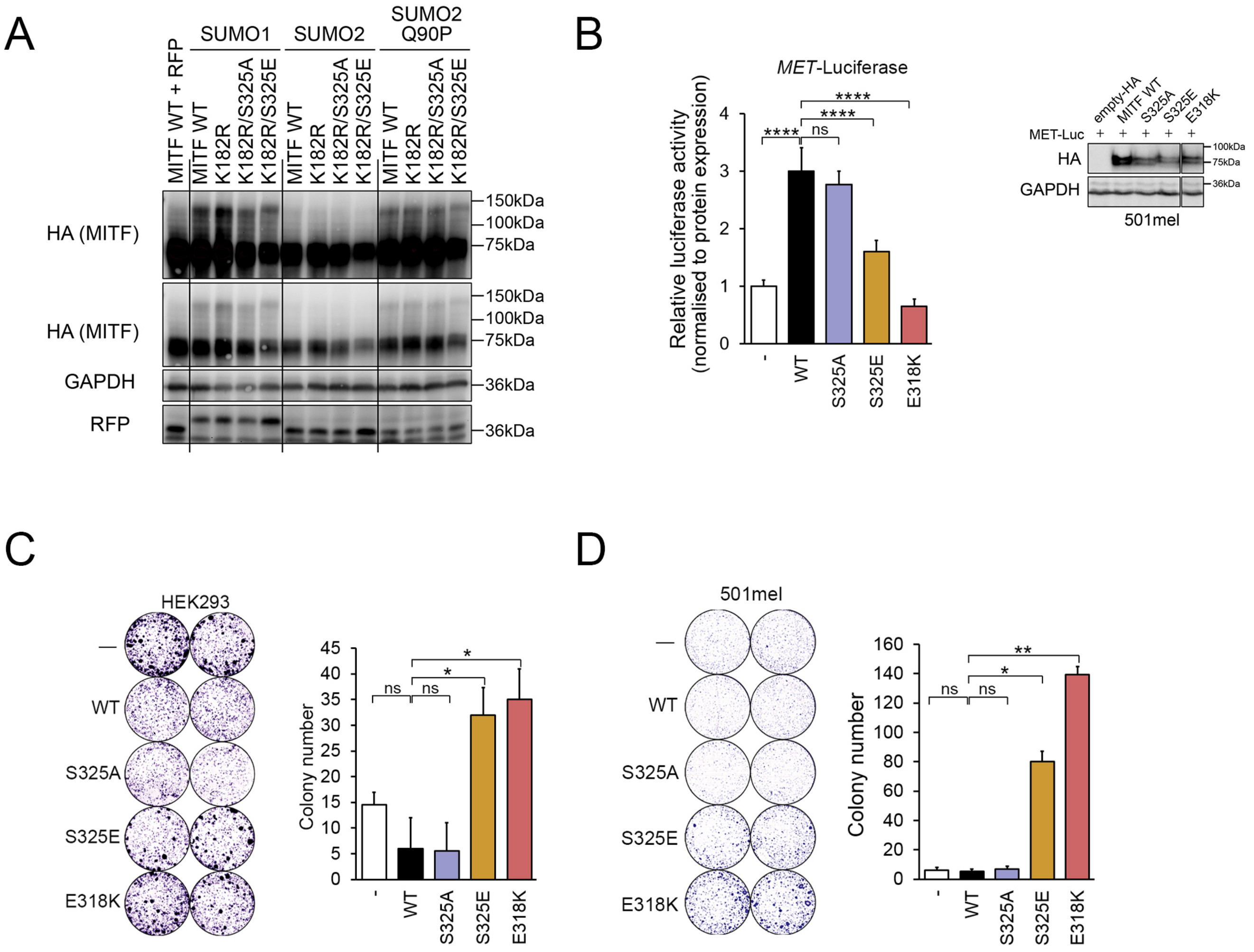


Binet et al. Supplemental Figure 4

**Figure S4.** (*A*) Western blot of HEK293 cells transiently co-transfected with various mutants of HA-MITF plus SUMO1-RFP, SUMO2-RFP or the non-cleavable SUMO2-Q90P mutant. GAPDH was used as loading control. (B) *Left*: Activity of the *MET* promoter when co-expressed with the indicated MITF mutants in 501mel cells. Luminescence is expressed in arbitrary values. Each mean was obtained from three independent transfections, each measured three time. Error bars: SD. The *p-values* were computed using the paired Wilcoxon Rank Sum test. *ns*: non-significant; ****: $p<0.0001$. *Right:* Western blot showing the relative expression of each MITF construct in the *Met* luciferase experiment. (*C*) Colony Formation Assay in HEK293 cells. Cells were fixed and stained with crystal violet, then scanned and colonies were counted. (*right*) Quantification from three independent transfections. Error bars: SD. The *p-values* were computed using the paired Wilcoxon Rank Sum test. *ns*: non-significant; *: $p<0.05$. (*D*) Colony Formation Assay in 501mel cells. Cells were fixed and stained with crystal violet, then scanned and colonies were counted. (*right*) Quantification from three independent transfections. Error bars: SD.

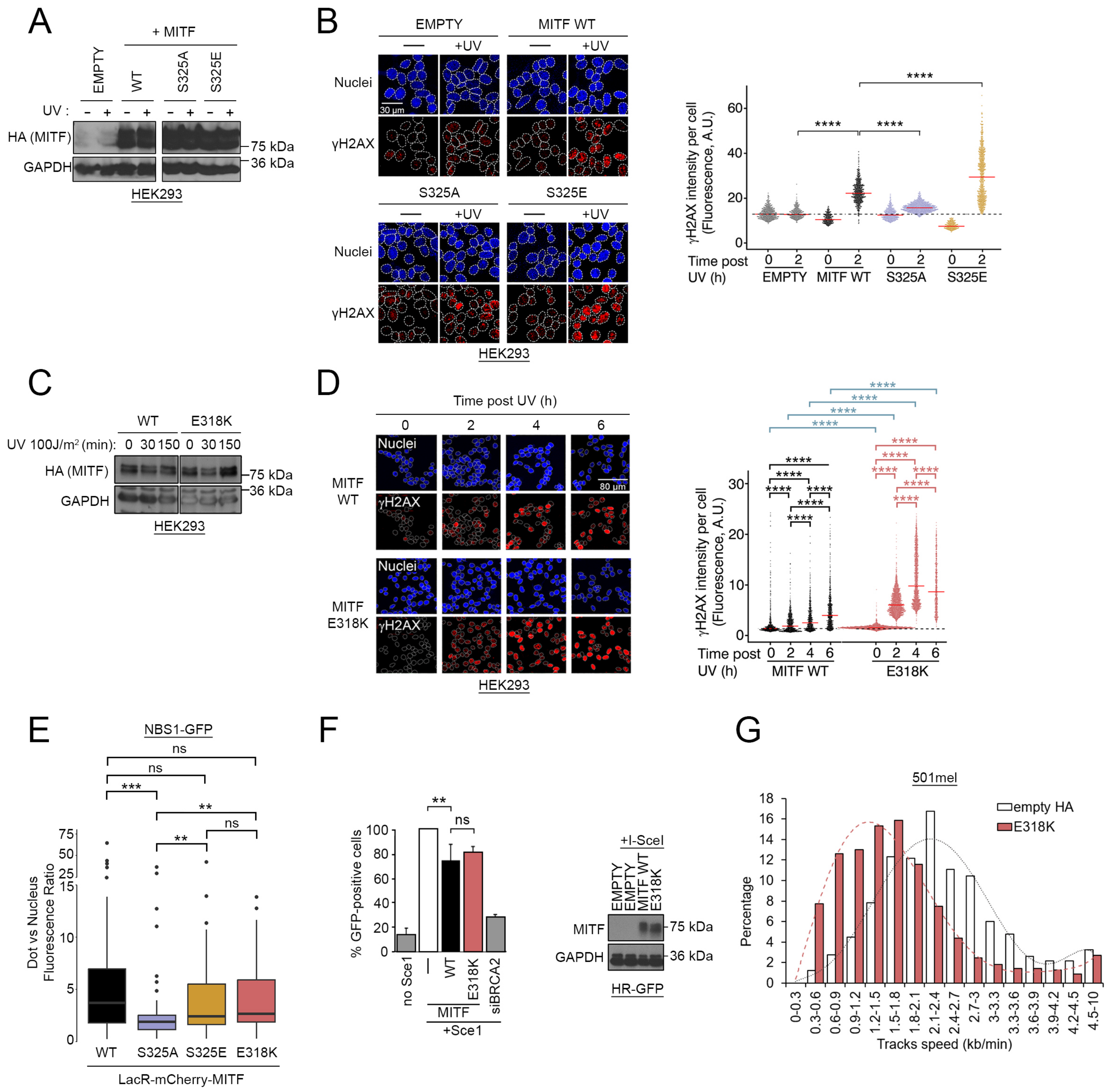


Binet et al. Supplemental Figure 5

**Figure S5.** (*A*) Western blot of HEK293 cells transfected with HA-tagged MITF WT, S325A or S325E mutants or the corresponding empty vector and exposed to UV (24 J/m$^2$). GAPDH was used a loading control. (*B*) Immunofluorescence analysis of γH2AX activation over time after UV irradiation (24 J/m$^2$). HEK293 cells were transfected with HA-MITF WT (black), S325A (purple) or S325E (gold) mutants or the corresponding empty vector (grey). Swarm plot representing the distribution of γH2AX intensities per cells. The medians are indicated in red. The black dotted line represents the median value of the EMPTY non-irradiated sample. (*C*) Western blot of HEK293 cells transfected with HA-tagged MITF WT or the E318K mutant and exposed to UV (100 J/m$^2$) for the indicated times. GAPDH was used a loading control. (*D*) Immunofluorescence analysis of γH2AX activation over time after UV irradiation (24 J/m$^2$). HEK293 cells were transfected with HA-MITF WT (black) or the E318K mutant (red). Swarm plot representing the distribution of γH2AX intensities per cells. The medians are indicated in red. The black dotted line represents the median value of the MITF WT non-irradiated sample. (*E*) Quantification of a nuclear tethering assay showing interaction between MITF WT or mutants and NBS1. The quantification is expressed as the ratio between the GFP fluorescence measured inside the area delimited by the mCherry dot and in the rest of the nucleus. The *p-values* were computed using the paired Wilcoxon Rank Sum test. *ns*: non-significant; **: $p<0.01$; ***: $p<0.001$. (*F*) *Left*: Graph showing the relative efficiency of homologous recombination using the U2-OS-DR-GFP reporter system. Data represent the mean (±SEM) from three independent experiments and are normalised against the I-SceI only samples. *Right*: Western blot of U2-OS-DR-GFP reporter cells transfected with MITF WT or the E318K mutant. GAPDH was used a loading control. (*G*) Histograms representing the distribution of the calculated forks speed in kilobase per minute in 501mel cells transfected with HA-MITF-E318K (red) or the corresponding empty vector (white). The dashed lines show the trendline for the matching bar graph.
